# G protein-coupled estrogen receptor activation by Bisphenol-A disrupts protection from apoptosis conferred by estrogen receptors ERα and ERβ in pancreatic beta cells

**DOI:** 10.1101/2022.01.31.478472

**Authors:** Ignacio Babiloni-Chust, Reinaldo S. dos Santos, Regla M. Medina-Gali, Atenea A. Perez-Serna, José-Antonio Encinar, Juan Martinez-Pinna, Jan-Ake Gustafsson, Laura Marroqui, Angel Nadal

## Abstract

17β-estradiol protects pancreatic β-cells from apoptosis via the estrogen receptors ERα, ERβ and GPER. Conversely, the endocrine disruptor Bisphenol-A (BPA), which exerts multiple effects in this cell type via the same estrogen receptors, increased basal apoptosis. The molecular initiated events that trigger these opposite actions have yet to be identified. We demonstrated that combined genetic downregulation and pharmacological blockade of each estrogen receptor increased apoptosis to a different extent. The increase in apoptosis induced by BPA was diminished by the pharmacological blockade or the genetic silencing of GPER, and it was partially reproduced by the GPER agonist G1. BPA and G1-induced apoptosis were abolished upon pharmacological inhibition, silencing of ERα and ERβ, or in dispersed islet cells from ERβ knockout (BERKO) mice. Yet, the ERα and ERβ agonists, PPT and DPN, respectively, had no effect on beta cell viability. To exert their biological actions, ERα and ERβ form homodimers and heterodimers. Molecular dynamic simulations together with proximity ligand assay and coimmunoprecipitation experiments indicated that the interaction of BPA with ERα and ERβ as well as the GPER activation by G1 decreased ERαβ heterodimers. We propose that ERαβ heterodimers play an antiapoptotic role in beta cells and that BPA- and G1-induced decrease in ERαβ heterodimers leads to beta cell apoptosis. Unveiling how different estrogenic chemicals affect the crosstalk among estrogen receptors should help to identify diabetogenic endocrine disruptors.

**Highlights:**

- Pharmacological blockade and gene silencing of estrogen receptors ERα, ERβ and GPER indicate that they are antiapoptotic in basal conditions.
- GPER activation by G1 and BPA triggered apoptosis via a crosstalk with ERα and ERβ.
- BPA interaction with ERα and ERβ as well as GPER activation decreased ERαβ heterodimers, which was associated to increased apoptosis.
- This pathway represents a novel molecular initiating event underlying the pro-apoptotic effect of BPA
- The EndoC-βH1 cell line may be a valid model of human β-cells for identifying diabetogenic pollutants.

## 1. Introduction

Loss of functional beta cell mass is a critical component contributing to the hyperglycemia observed in individuals with type 1 or type 2 diabetes. The interaction between the individual genetic background and environmental factors determines the progression of beta cell dysfunction and death (Eizirik et al., 2020).

Sex differences exist in the prevalence of diabetes. Premenopausal women have a lower incidence of diabetes compared with men (Gannon et al., 2018; Kautzky-Willer et al., 2016). These sex differences can be partially attributed to the actions of 17β-estradiol (E2) through three estrogen receptors (ERs), namely estrogen receptors α (ERα) and β (ERβ) (Heldring et al., 2007) and the G protein-coupled estrogen receptor (GPER/GPR30) (Revankar et al., 2005; Thomas et al., 2005). After ligand binding, ERα and ERβ can either form homo- and heterodimers and act as transcription factors or tether other DNA-bound transcription factors (Heldring et al., 2007). In addition to their nuclear-initiated actions, ERα and ERβ can trigger extranuclear-initiated effects after dimerization and activate a variety of signaling pathways (Levin and Hammes, 2016).

All three ERs protect beta cells from different apoptotic insults (Balhuizen et al., 2010; le May et al., 2006; Liu et al., 2009), yet the role of E2 can be disrupted by environmental chemicals that compete for the same receptors and modify the physiological pathways activated by the natural hormone (Gore et al., 2015). Bisphenol-A (BPA), which is used as a model of endocrine disrupting chemical that alters insulin secretion and insulin sensitivity in mice (Alonso-Magdalena et al., 2010, 2006), has been implicated in the etiology of diabetes (Alonso-Magdalena et al., 2011; Sargis and Simmons, 2019). At concentrations similar to those found in human blood (Vandenberg et al., 2010), BPA increases pancreatic insulin content via ERα (Alonso-Magdalena et al., 2008) and regulates ion channel expression and function as well as augments insulin release in an ERβ-mediated manner (Marroqui et al., 2021; Martinez-Pinna et al., 2019; Soriano et al., 2012). In INS-1 cells and mouse dispersed islet cells, BPA induces mitochondrial dysfunction, ROS production and NF-κB activation, which culminates in beta cell apoptosis (Carchia et al., 2015; Lin et al., 2013).

As E2 and BPA have opposite actions on beta cell survival, we decided to investigate the molecular initiating event (MIE) (Allen et al., 2014) underlying the pro-apoptotic effect of BPA.

Using several beta cell models, including the EndoC-βH1 cell line, which is a valid model of human beta cells recommended for screening studies (Tsonkova et al., 2018), we show that ERα, ERβ and GPER mediate beta cell survival. Our findings suggest that ERαβ heterodimerization is key to this antiapoptotic role and that G1, a GPER agonist, and BPA induce apoptosis via a mechanism involving the reduction in ERαβ heterodimers downstream of GPER.

## 2. Material and methods

### 2.1. Chemical substances and animals

Bisphenol-A was purchased from MP Biomedicals (Cat no. 155118; Santa Ana, CA, USA). 17β-estradiol (E2, Cat no. E8875) was obtained from Sigma-Aldrich (Saint Louis, MO, USA). Propylpyrazoletriol (PPT, Cat no. 1426), Diarylpropionitrile (DPN, Cat no. 1494), G1 (Cat no. 3577), ICI 182,780 (Cat no. 1047), PHTPP (Cat no. 2662) and Methylpiperidinopyrazole (MPP, Cat no. 1991) were obtained from Tocris Cookson (Bristol, UK).

Mice with knockout of the *Erβ* gene (also known as *Esr2*) (BERKO mice), supplied by Jan-Åke Gustafsson’s laboratory, were generated as described (Krege et al., 1998). Wild-type littermates and BERKO mice were obtained from the same supplier and colony and kept under standard housing conditions (12 h light/dark cycle, food *ad libitum*). Experimental procedures were performed according to the Spanish Royal Decree 1201/2005 and the European Community Council directive 2010/63/EU. The ethical committee of Miguel Hernandez University reviewed and approved the methods used herein (approvals ID: UMH-IB-AN-01–14 and UMH-IB-AN-02-14).

### 2.2. Culture of cell lines and dispersed islet cells

Rat insulin-producing INS-1E cells (RRID: CVCL_0351, kindly provided by Dr. C. Wollheim, Department of Cell Physiology and Metabolism, University of Geneva, Geneva, Switzerland) were cultured as previously described (Santin et al., 2016). Human insulin-producing EndoC-βH1 cells (RRID: CVCL_L909, Univercell-Biosolutions, France) were cultured in Matrigel/fibronectin-coated plates as described before (Ravassard et al., 2011). INS-1E and EndoC-βH1 cells have been shown to be free of mycoplasma infection. Pancreatic islets were isolated using collagenase (Sigma, St Louis, MO, USA) as described (Alonso-Magdalena et al., 2008). Islets were dispersed into single cells and cultured in polylysine-coated plates as described before (Martinez-Pinna et al., 2019). Cell lines and dispersed cells were kept at 37°C in a humidified atmosphere of 95% O_2_ and 5% CO_2_.

### 2.3. RNA interference

Optimal siRNA concentration (30 nM) and conditions for small interfering RNA (siRNA) transfection using Lipofectamine RNAiMAX lipid reagent (Invitrogen, Carlsbad, CA, USA) were previously established (Santin et al., 2016). Allstars Negative Control siRNA (Qiagen, Venlo, the Netherlands) was used as a negative control (siCTRL). siRNA targeting *Erα* (also known as *Esr1*; si*ERα*), *Erβ* (si*ERβ*) or *Gper1* (si*GPER1*) (Qiagen, Venlo, the Netherlands) were used herein (Supplementary Table 2). Of note, GPER silencing in INS-1E cells as well as ERα and ERβ silencing in EndoC-βH1 cells were performed in a two-step transfection protocol (Marroqui et al., 2015). Briefly, cells were exposed to 30 nmol/l siCTRL or si*GPER1*/si*Gper1* for 16 h, washed and allowed to recover in culture for 24 h. Next, cells were exposed again to the same siRNAs for 16 h, allowed to recover in culture for 48 h, and then used for the subsequent experiments. Transfections with siRNAs were carried out in BSA- and antibiotic-free medium.

### 2.4. Cell viability assessment by DNA-binding dyes

Percentage of apoptosis was determined after staining with the DNA-binding dyes Hoechst 33342 and propidium iodide as described (Santin et al., 2016). To avoid bias, cell viability was assessed by two different observers, being one of them unaware of sample identity. The agreement of results between both observes was higher than 90%.

### 2.5. Flow cytometric analysis

Apoptotic cells were analyzed by flow cytometry using the FITC Annexin V Apoptosis Detection Kit with propidium iodide (BioLegend, San Diego, CA, USA) following the manufacturer’s instructions. Briefly, INS-1E cells were detached and dissociated using Accutase (Thermo Fisher Scientific). Cell suspension was washed twice with PBS (200 *g* for 7 min) and proceeded to Annexin V-fluorescein isothiocyanate (FITC)/PI staining according to the manufacturer’s instructions (FITC Annexin V Apoptosis Detection Kit with PI; BioLegend, San Diego, CA, USA). Cells stained with Annexin V (both Annexin V^+^- and Annexin V^+^/PI^+^-cells) were detected using a FACSCanto II (BD Biosciences, Madrid, Spain) flow cytometer.

### 2.6. Caspase 3/7 activity

Caspase 3/7 activity was determined using the Caspase-Glo® 3/7 assay (Promega, Madison, WI, USA) following the manufacturer’s instructions. Briefly, following treatment, cells were incubated with Caspase-Glo® 3/7 reagent at room temperature before recording luminescence with a POLASTAR plate reader (BMG Labtech, Germany).

### 2.7. MTT assay

Cell viability was measured by the colorimetric assay showing reduction of MTT (3-(4,5-dimethylthiazol-2-yl)-2,5-diphenyltetrazolium bromide) (Sigma-Aldrich) as previously described (Denizot and Lang, 1986; Mosmann, 1983). Briefly, MTT prepared in RPMI 1640 without phenol red was added (final concentration: 0.5 mg/ml) and incubated at 37°C for 3 h. Upon incubation, the supernatant was aspirated and 100 ml DMSO was added to dissolve formazan crystals. The absorbance was measured at 595 nm using an iMark™ Microplate absorbance reader (Bio-Rad, Hercules, CA) and the percentage of cell viability was calculated.

### 2.8. Real-time PCR

Quantitative RT-PCR was carried out in a CFX96 Real-Time System (Bio-Rad Laboratories, Hercules, CA, USA). RNA was extracted with the RNeasy Micro kit (Qiagen) and cDNA prepared with the High-Capacity cDNA Reverse Transcription kit (Applied Biosystems, Foster City, CA, USA). Amplification reactions were performed as describe (Villar-Pazos et al., 2017). Values were analyzed with CFX Manager Version 1.6 (Bio-Rad) and expressed as the relative expression in respect of control values (2^−ΔΔCt^) (Schmittgen and Livak, 2008). *Gapdh* and β-actin were used as housekeeping genes for rat and human samples, respectively. The primers used herein are listed in Supplementary Table 2.

### 2.9. DCF assay

Oxidative stress was measured using the fluorescent probe 2’,7’-dichlorofluorescein diacetate (DCF; Sigma-Aldrich) as described (Cunha et al., 2016). Briefly, cells seeded in 96-well black plates were loaded with 10 μM DCF for 30 min at 37°C and washed with PBS. DCF fluorescence was quantified in a POLASTAR plate reader (BMG Labtech, Germany). Data are expressed as DCF fluorescence corrected by total protein.

### 2.10. Western blotting

Cells were washed with cold PBS and lysed in Laemmli buffer. Immunoblot analysis was performed by overnight incubation with the antibodies against ERα, ERβ, GPER, β-actin, GAPDH and α-tubulin. Afterwards, membranes were incubated for 1 h at room temperature with peroxidase-conjugated antibodies (1:5000) as secondary antibodies. SuperSignal West Femto chemiluminescent substrate (Thermo Scientific, Rockford, IL, USA) and ChemiDoc XRS+ (Bio-Rad Laboratories) were used to detect immunoreactive bands. Densitometry analysis was done with the Image Lab software (version 4.1, Bio-Rad Laboratories). The antibodies used herein are listed in Supplementary Table 3.

### 2.11. In situ proximity ligation assay (PLA)

In situ PLA was performed using a Duolink® In Situ Red Starter Kit (Sigma-Aldrich) following the manufacturer’s instructions with slight modifications (INCLUDE Iwabuchi E, Miki Y, Ono K, et al, J Steroid Biochem Mol Biol 165:159–169, 2017)(Iwabuchi et al., 2017). Briefly, cells grown on coverslips were washed with PBS and fixed with 4% paraformaldehyde (VWR Chemicals, Spain). Then, cells were washed with PBS and permeabilized with 0.1% Triton-X-100. Subsequently, cells were incubated for 30 min at 37ºC with a blocking solution and incubated overnight at 4ºC with the corresponding primary antibodies (Supplementary Table 3). PLA probe solution was added and incubated for 1 h at 37ºC. Ligation-Ligase solution was added, and coverslips were incubated for 30 min at 37ºC. Subsequently, amplification-polymerase solution was added and incubated for 100 min at 37ºC. Of note, all incubations at 37ºC were carried out using a pre-heated humidity chamber. Finally, coverslips were washed with a specific buffer provided and mounted with a minimal volume of Duolink in situ Mounting Medium with DAPI®. ER dimers were observed using a Zeiss Confocal LSM900 microscope equipped with a camera (Zeiss-Vision, Munich, Germany) and images were acquired at x63 magnification and analyzed using ZEN software (version 3.2; Zeiss-Vision, Munich, Germany).

### 2.12. Co-immunoprecipitation

INS-1E cells were washed with cold PBS, lysed in cold immunoprecipitation buffer (50 mmol/l Tris, pH 7.5, 4 mmol/l NaCl, 2 mmol/l MgCl_2_, 10 mmol/l NaF, 1 mmol/l PMSF, 1% Triton X-100 and Complete Protease inhibitor mixture, Roche Diagnostics) for 30 min on ice and centrifuged at 20,000 *g* for 10 min at 4ºC. Cell lysates were pre-cleared for 1 h at 4ºC with Dynabeads Protein G (Thermo Fischer Scientific; Cat no. 10003D). The same amounts of protein were incubated overnight at 4ºC, either with an anti-ERβ antibody or nonspecific rabbit IgG (Santa Cruz Biotechnology) used as negative control. Upon overnight incubation, Immunoprecipitates were incubated for 1 h at 4ºC with Dynabeads Protein G, washed six times with cold immunoprecipitation buffer and resuspended in 5x Laemmli buffer. Immunoprecipitates and total protein (Input) were subjected to SDS-PAGE and immunoblotted with mouse anti-ERα antibody. The antibodies used herein are listed in Supplementary Table 3.

### 2.13. Molecular docking and dynamics simulations

More than 300 human ERα-ligand-binding domain (LBD) (UniProt code: P03372) and 32 human ERβ-LBD (UniProt code: Q92731) structures were resolved from cryptographic data. As these structures contained unresolved residues in mobile regions of the protein that diffracted poorly, the lost amino acids were reconstructed after generation of a homology model at the Swiss-Model server (Biasini et al., 2014; Marroqui et al., 2021). After unresolved gaps elimination, models for the hERα-LBD monomer (Protein Data Bank entry 5DXE as template) and the hERβ-LBD monomer (Protein Data Bank entry 3OLS as template) were generated. From these monomeric structures homo- and heterodimers were reconstructed using GRAMM-X server (Tovchigrechko and Vakser, 2006). E2 and BPA molecular docking simulations within the cavity of each LBD were performed using the YASARA structure software (version 20.12.24) (Marroqui et al., 2021). Molecular dynamics simulations on the structures with the best docking calculations were performed with YASARA structure software (version 20.10.24) (Marroqui et al., 2021). The intermolecular protein interaction energy for homo- and heterodimers subunits was calculated using the Folds 5.0 software (Delgado et al., 2019).

### 2.14. Statistical analysis

Experimenters were not blind to group assignment and outcome assessment. GraphPad Prism 7.0 software (GraphPad Software, La Jolla, CA, USA; www.graphpad.com) was used for all statistical analyses. Data are expressed as the mean ± SEM. To assess differences between groups, we used two-tailed Student’s *t* test or ANOVA when appropriate. For nonparametric data we used Mann–Whitney and Kruskal–Wallis ANOVA tests (followed by Dunn’s test), depending on the experimental groups involved in the comparison. A *p* value ≤0.05 was considered statistically significant.

## 3. Results

In INS-1E (Fig. 1a) and EndoC-βH1 cells (Fig. 1b), E2 (10 pM/l to 1 µM) either had no effect or decreased apoptosis. Conversely, BPA increased apoptosis in both cell lines in a dose-dependent manner (Fig. 1c,d). Similar results were observed in dispersed islet cells from female mice (Fig. 1e). These results were confirmed by three other approaches, namely flow cytometry analysis (Fig. 1f), caspase 3/7 activity (Fig. 1g) and MTT assay (Supplementary Fig. 1). When E2 and BPA were added together, E2 1 nM prevented BPA-induced apoptosis (Fig. 1h), suggesting that E2 and BPA initiate common pathways. Previous work demonstrated that BPA induces oxidative stress in beta cells (Carchia et al., 2015; Lin et al., 2013). Similarly, we observed that BPA upregulated genes encoding the antioxidant enzymes, superoxide dismutase (*Sod2*), glutathione peroxidase 4 (*Gpx4*) and catalase (*Cat*) (Supplementary Fig. 2a-c) and increased reactive oxygen species (ROS) generation (Supplementary Fig. 2d,e). The antioxidant N-acetylcysteine abolished BPA-induced ROS production in INS-1E (Supplementary Fig. 2d) and EndoC-βH1 cells (Supplementary Fig. 2e). Notably, N-acetylcysteine prevented BPA-induced apoptosis in both cell lines (Supplementary Fig. 2f,g), reinforcing that oxidative stress is involved in BPA-induced apoptosis. Of note, E2 did not change ROS levels (data not shown). These results indicate that ROS production is a key event involved in BPA-induced beta cell apoptosis.

**Fig. 1.**
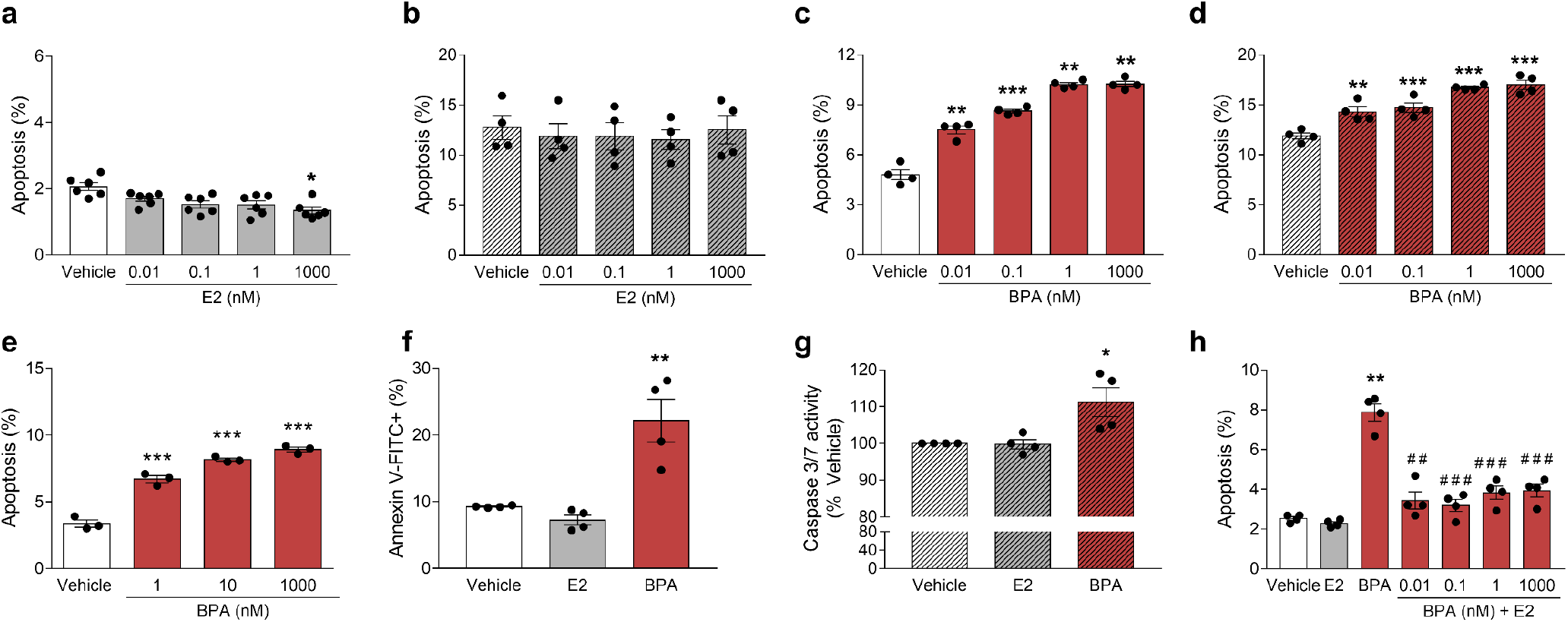
E2 and BPA have different effects on beta cell viability. INS-1E (**a**,**c**) and EndoC-βH1 cells (**b**,**d**) were treated with vehicle (white bars), E2 (grey bars) or BPA (red bars) for 24 h. (**e**) Dispersed islet cells from female mice were treated with vehicle (white bars) or BPA (red bars) for 48 h. Apoptosis was evaluated using Hoechst 33342 and propidium iodide staining. (**f**) INS-1E cells were treated with vehicle (white bar), E2 1 nM(grey bar) or BPA 1 nM (red bar) for 48 h. Annexin V-FITC-positive cells were analyzed by flow cytometry. (**g**) EndoC-βH1 cells were treated with vehicle (white bar), E2 1 nM (grey bar) or BPA 1 nM (red bar) for 48 h. Caspase 3/7 activity was measured by a luminescent assay. Results are expressed as % vehicle-treated cells. (**h**) INS-1E were treated with vehicle (white bar), E2 1 nM (grey bar), BPA 1 nM (red bar) or a combination of both (red bars + E2) for 24 h. Apoptosis was evaluated using Hoechst 33342 and propidium iodide staining. Data are shown as means ± SEM of 3-6 independent experiments. (**a-i**) \**p*≤0.05, \*\**p*≤0.01 and \*\*\**p*≤0.001 vs Vehicle, by one-way ANOVA. (**h**) \*\**p*≤0.01 vs Vehicle; ^##^*p*≤0.01 and ^###^*p*≤0.001 vs BPA 1 nM. One-way ANOVA.

As previous results indicate that BPA acts via ERs in beta cells (Alonso-Magdalena et al., 2008; Martinez-Pinna et al., 2019; Soriano et al., 2012) we evaluated the expression of ERα, ERβ and GPER in INS-1E and EndoC-βH1 cells by quantitative RT-PCR and western blot. Considering ERα expression as 1, mRNA analyses showed that the ratio GPER:ERα:ERβ was 100:1:0.3 for INS-1E (Supplementary Fig. 3a) and 675:1:3.3 for EndoC-βH1 cells (Supplementary Fig. 3b). This expression pattern was confirmed at protein level, where GPER was the most expressed of the three receptors (Supplementary Fig. 3c).

### 3.1. GPER mediates BPA-induced apoptosis

We first assessed whether GPER participated in BPA-induced apoptosis. GPER antagonist G15 reduced BPA-triggered apoptosis in INS-1E and EndoC-βH1 cells (Fig. 2a,b). Doses as low as 100 pM of the GPER agonist G1 induced apoptosis in INS-1E cells (Supplementary Fig. 4a). In comparison with BPA 1 nM, G1 100 nM presented a significantly smaller effect on apoptosis in both cell lines (Fig. 2c,d). To further study the GPER role in beta cell survival, we used siRNAs to inhibit GPER expression in INS-1E and EndoC-βH1 cells (Fig. 2e-j, Supplementary Fig. 4b,c). GPER silencing promoted apoptosis in basal conditions in INS-1E but not in EndoC-βH1 cells (Fig. 2f,i). Moreover, GPER-inhibited cells were less susceptible to BPA-induced apoptosis than control cells (Fig. 2f,g,i,j). Similar data were obtained with a second, independent siRNA (data not shown). These results indicate that GPER activation is part of the mechanism whereby BPA elicits apoptosis.

**Fig. 2.**
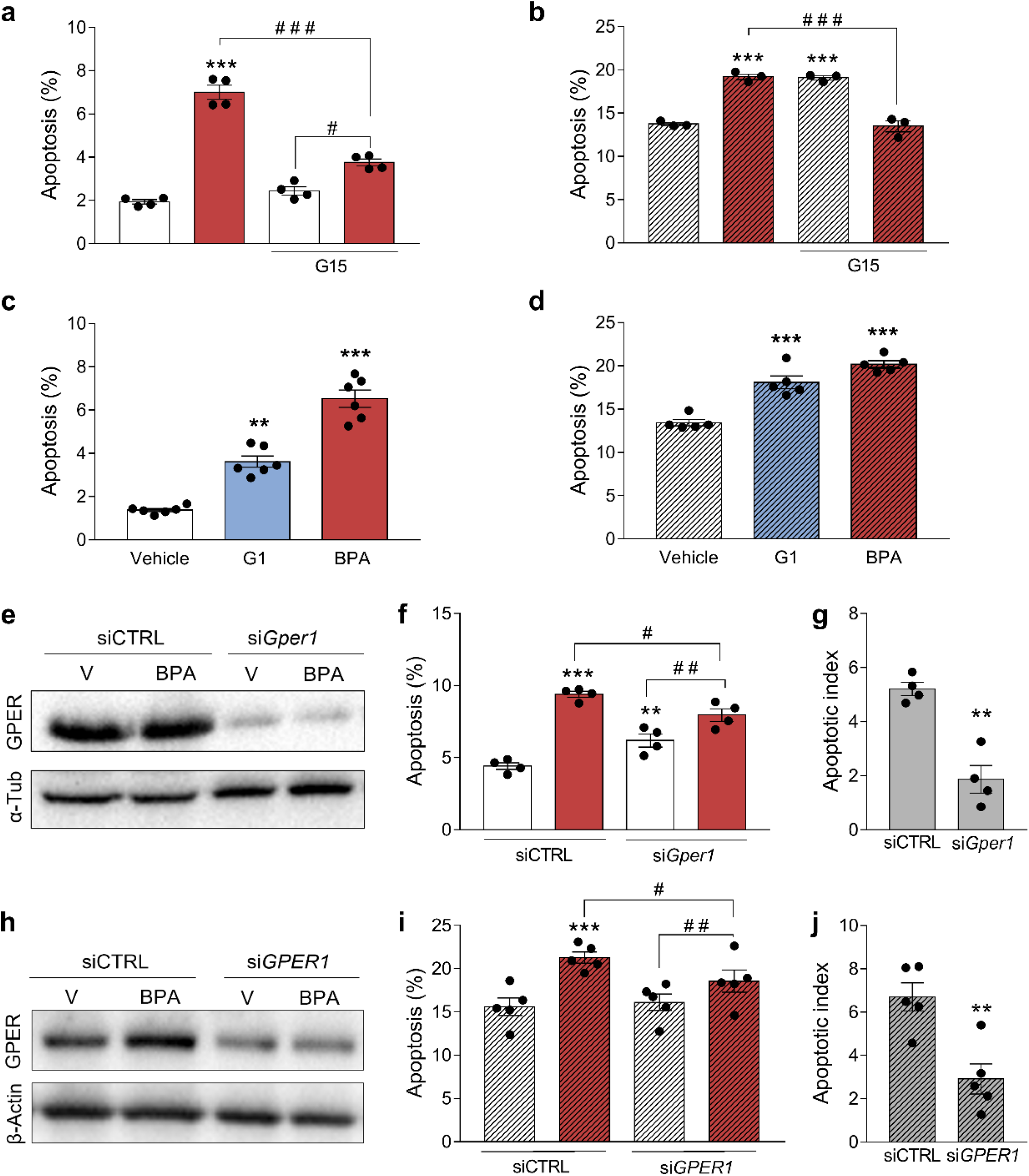
BPA-induced apoptosis requires GPER. (**a**,**b**) INS-1E (**a**) and EndoC-βH1 cells (**b**) were pre-treated with vehicle or GPER antagonist G15 10 nM for 3 h. Afterwards, cells were treated with vehicle (white bars) or BPA 1 nM (red bars) in the absence or presence of G15 10 nM for 24 h. (**c**,**d**) INS-1E (**c**) and EndoC-βH1 cells (**d**) were treated with vehicle (white bars), G1 100 nM (blue bars) or BPA 1 nM (red bars) for 24 h. Apoptosis was evaluated using Hoechst 33342 and propidium iodide staining. (**e-j**) INS-1E (**e-g**) and EndoC-βH1 cells (**h-j**) were transfected with siCTRL or with a siRNA targeting GPER (si*Gper1* or si*GPER1*). Cells were treated with vehicle (white bars) or BPA 1 nM (red bars) for 24 h. (**e**,**h**) Protein expression was measured by western blot. Representative images of four (**e**) or five (**h**) independent experiments are shown. (**f**,**i**) Apoptosis was evaluated using Hoechst 33342 and propidium iodide staining. (**g**,**j**) BPA-induced apoptosis data from Fig. 5f (**g**) and Fig. 5i (**j**) are presented as apoptotic index. Data are shown as means ± SEM of 4-6 independent experiments. (**a**,**b**) \*\**p*≤0.01 and \*\*\**p*≤0.001 vs untreated vehicle; ^##^*p*≤0.01, ^###^*p*≤0.001 as indicated by bars. Two-way ANOVA. (**c**,**d**) \*\**p*≤0.01 and \*\*\**p*≤0.001 vs vehicle, by one-way ANOVA. (**f**,**i**) \*\**p*≤0.01 and \*\*\**p*≤0.001 vs siCTRL vehicle; ^#^*p*≤0.05 and ^##^*p*≤0.01 as indicated by bars. Two-way ANOVA. (**g**,**j**) \*\**p*≤0.01 vs siCTRL, by two-tailed Student’s *t* test.

### 3.2. ERα and ERβ are involved in BPA-induced apoptosis

We then investigated whether ERα and ERβ were also implicated in BPA-induced apoptosis. In both cell lines, the pure antiestrogen ICI 182,780 abolished the BPA effect on apoptosis (Fig. 3a,b). Treatment with the ERα antagonist methylpiperidinopyrazole (MPP) induced apoptosis in basal conditions, while propylpyrazoletriol (PPT) an ERα agonist, did not change viability (Fig. 3c, Supplementary Fig. 5a,b). In INS-1E cells, MPP abolished BPA-induced apoptosis (Fig. 3c). To further characterize ERα participation in cell survival and BPA-induced apoptosis, we silenced ERα expression in INS-1E and EndoC-βH1 cells using specific siRNAs (Fig. 3d,g, Supplementary Fig. 5c-e). ERα knockdown augmented apoptosis in basal conditions and prevented BPA-induced apoptosis in both cell lines (Fig. 3e,f,h,i). We confirmed these data with a second, independent siRNA (data not shown).

**Fig. 3.**
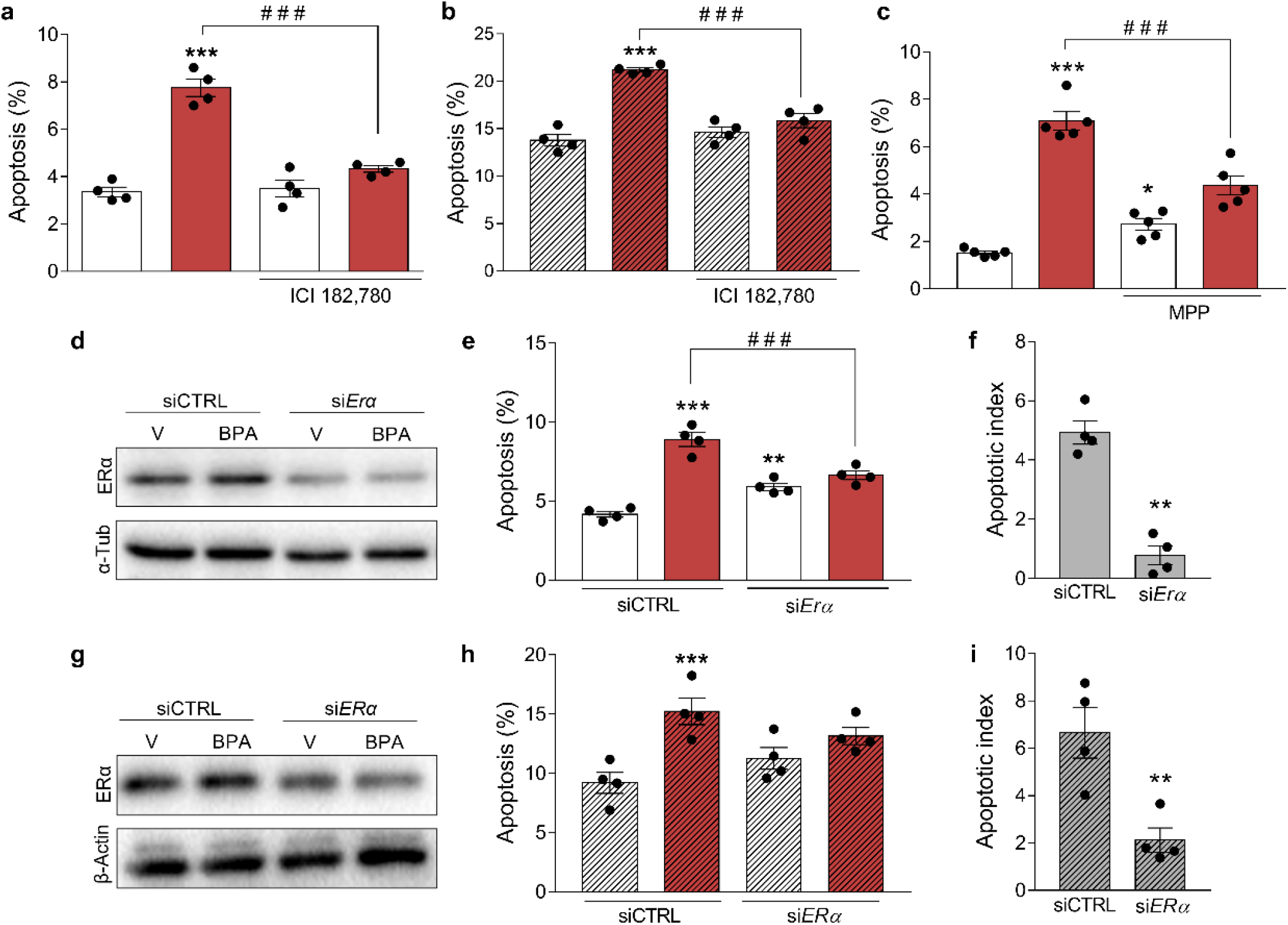
ERα mediates BPA-induced apoptosis. (**a**,**b**) INS-1E (**a**) and EndoC-βH1 cells (**b**) were pre-treated with vehicle or antiestrogen ICI 182,780 1 µM for 3 h. Afterwards, cells were treated with vehicle (white bars) or BPA 1 nM (red bars) in the absence or presence of ICI 182,780 1 µM for 24 h. (**c**) INS-1E cells were pre-treated with vehicle or ERα antagonist MPP 100 nM for 3 h. Afterwards, cells were treated with vehicle (white bars) or BPA 1 nM (red bars) in the absence or presence of MPP 100 nM for 24 h. Apoptosis was evaluated using Hoechst 33342 and propidium iodide staining. (**d-i**) INS-1E (**d-f**) and EndoC-βH1 cells (**g-i**) were transfected with siCTRL or with a siRNA targeting ERα (si*Erα* or si*ERα*). Cells were treated with vehicle (white bars) or BPA 1 nM (red bars) for 24 h. (**d**,**g**) Protein expression was measured by western blot. Representative images of four independent experiments are shown. (**e**,**h**) Apoptosis was evaluated using Hoechst 33342 and propidium iodide staining. (**g**,**j**) BPA-induced apoptosis data from Fig. 5e (**f**) and Fig. 5h (**i**) are presented as apoptotic index. Data are shown as means ± SEM of 4-5 independent experiments. (**a-c**) \**p*≤0.05 and \*\*\**p*≤0.001 vs untreated vehicle; ^###^*p*≤0.001 as indicated by bars. Two-way ANOVA. (**e**,**h**) \*\**p*≤0.01 and \*\*\**p*≤0.001 vs siCTRL vehicle; ^###^*p*≤0.001 as indicated by bars. Two-way ANOVA. (**f**,**i**) \*\**p*≤0.01 vs siCTRL, by two-tailed Student’s *t* test.

Regarding ERβ, while its antagonist 4-(2-phenyl-5,7-bis(trifluoromethyl)pyrazolo[1,5-a]pyrimidin-3-yl)phenol (PHTPP) also induced apoptosis in basal conditions, ERβ agonist diarylpropionitrile (DPN) had no effect on beta cell viability (Fig. 4a, Supplementary Fig. 6a,b). ERβ inhibition by siRNAs (Fig. 4c,f, Supplementary Fig. 6c-h) induced a substantial increase in apoptosis, mainly in INS-1E cells; importantly, BPA-elicited apoptosis was partially lost following ERβ silencing (Fig. 4d,e,g,h). Comparable results were observed with a second, independent siRNA (data not shown). Finally, BPA effects on viability were abrogated in dispersed islets cells from BERKO mice (Fig. 4b).

**Fig. 4.**
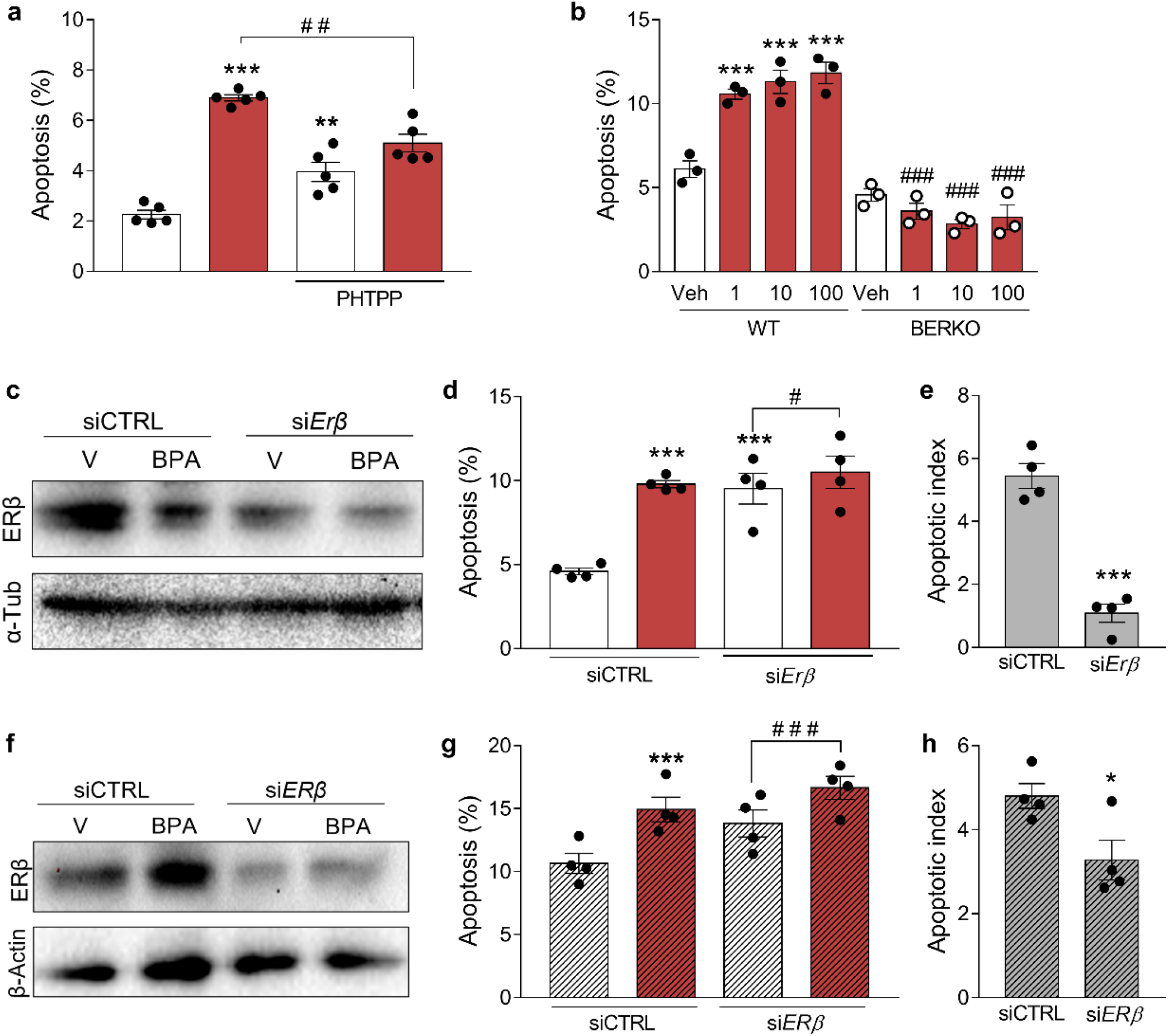
ERβ mediates BPA-induced apoptosis. (**a**) INS-1E cells were pre-treated with vehicle or ERβ antagonist PHTPP 1 µM for 3 h. Afterwards, cells were treated with vehicle (white bars) or BPA 1 nM (red bars) in the absence or presence of PHTPP 1 µM for 24 h. (**b**) Dispersed islet cells from WT and BERKO mice were treated with vehicle (white bars) or BPA (red bars) for 48 h. Apoptosis was evaluated using Hoechst 33342 and propidium iodide staining. (**c-h**) INS-1E (**c-e**) and EndoC-βH1 cells (**f-h**) were transfected with siCTRL or with a siRNA targeting ERβ (si*Erβ* or si*ERβ*). Cells were treated with vehicle (white bars) or BPA 1 nM (red bars) for 24 h. (**c**,**f**) Protein expression was measured by western blot. Representative images of four independent experiments are shown. (**d**,**g**) Apoptosis was evaluated using Hoechst 33342 and propidium iodide staining. (**e**,**h**) BPA-induced apoptosis data from Fig. 5d (**e**) and Fig. 5g (**h**) are presented as apoptotic index. Data are shown as means ± SEM of 3-5 independent experiments. (**a**) \*\**p*≤0.01 and \*\*\**p*≤0.001 vs untreated vehicle; ^##^*p*≤0.01 as indicated by bars. Two-way ANOVA. (**b**) \*\*\**p*≤0.001 vs WT vehicle; ^###^*p*≤0.001 vs respective WT. Two-way ANOVA. (**d**,**g**) \*\*\**p*≤0.001 vs siCTRL vehicle; ^#^*p*≤0.05 and ^###^*p*≤0.001 as indicated by bars. Two-way ANOVA. (**e**,**h**) \**p*≤0.05 and \*\*\**p*≤0.001 vs siCTRL, by two-tailed Student’s *t* test.

### 3.3. A crosstalk among ERα, ERβ and GPER mediates beta cell survival

As our data indicate that GPER signaling triggers beta cell apoptosis via downstream activation of ERα and ERβ, we sought to examine this hypothesis. First, we observed that G1-induced apoptosis was abolished by the antiestrogen ICI 182,780 (Fig. 5a,b). As G1 may also bind to and activate a 36-kDa variant of ERα, ERα36 (Kang et al., 2010), we silenced GPER (Fig. 5c, Supplementary Fig. 7a) to test whether G1 affected apoptosis via GPER. Notably, G1 did not affect apoptosis in GPER-silenced cells (Fig. 5d, Supplementary Fig. 7b), indicating that, in our cell system, G1 acts preferentially through GPER. Moreover, MPP (ERα antagonist) and PHTPP (ERβ antagonist) prevented G1-induced apoptosis (Fig. 5e-h). We then tested G1 effects on apoptosis upon silencing of ERα or ERβ separately (si*Erα* and si*Erβ*) or simultaneously (si*Erα*/*Erβ*) (Fig. 5i, Supplementary Fig. 7c-e). As described above, ERα or ERβ knockdown induced beta cell death and silencing of both receptors simultaneously induced apoptosis to the same extent observed in ERβ-inhibited cells (Fig. 5i, Supplementary Fig. 7f). G1 effect on apoptosis was lost in ERα-, ERβ-or ERα/ERβ-deficient cells (Fig. 5i, Supplementary Fig. 7f). Lastly, G1 induced apoptosis in dispersed islet cells from WT mice but its effect on apoptosis was completely blunted in dispersed cells from BERKO mice (Fig. 5j). These results indicate that ERβ seems to be more important than ERα for beta cell survival and that ERα and ERβ are required for GPER effects on viability.

**Fig. 5.**
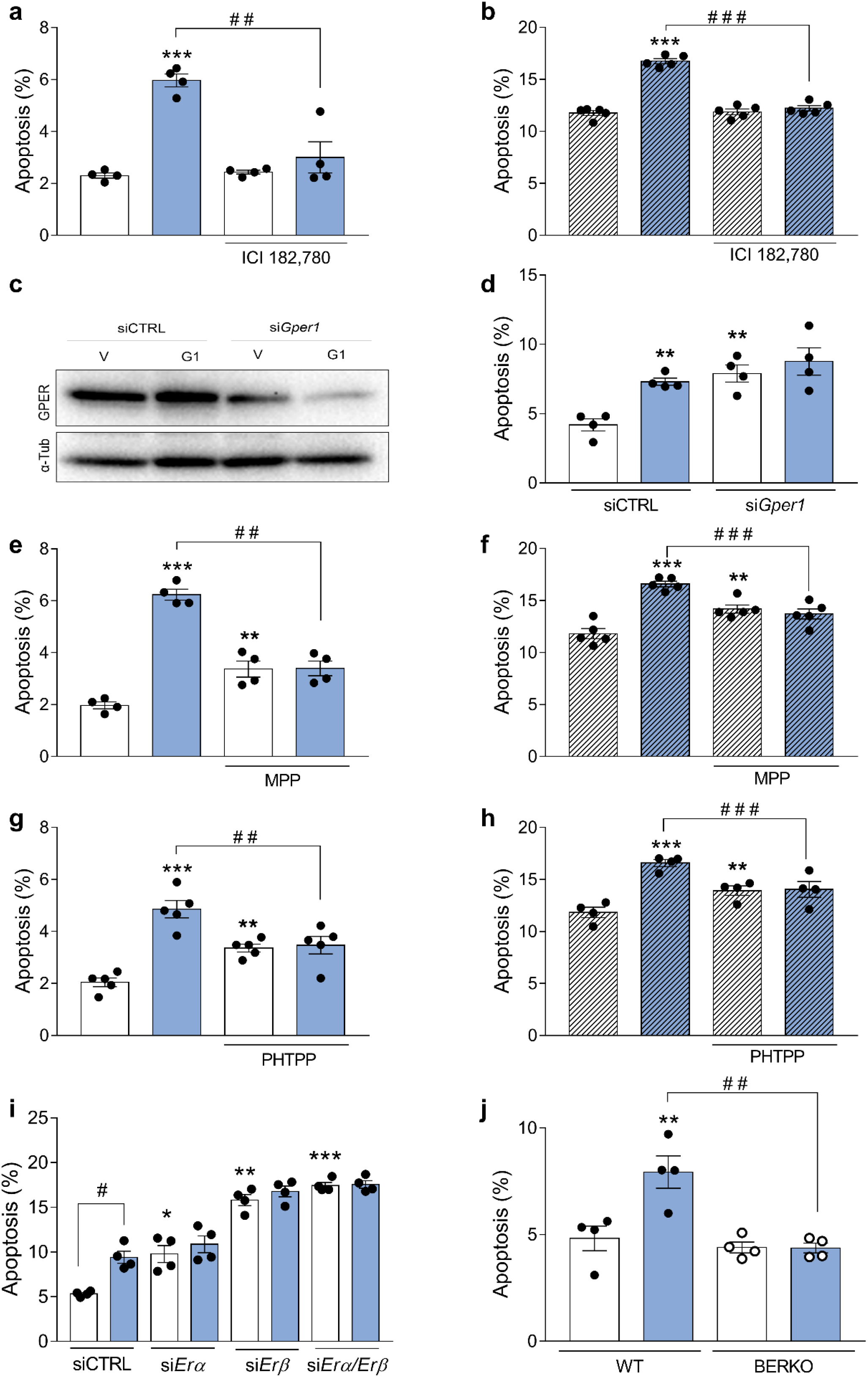
A GPER crosstalk with ERα and ERβ induces apoptosis. (**a**,**b**) INS-1E (**a**) and EndoC-βH1 cells (**b**) were pre-treated with vehicle or antiestrogen ICI 182,780 1 µM for 3 h. Afterwards, cells were treated with vehicle (white bars) or G1 100 nM (blue bars) in the absence or presence of ICI 182,780 1 µM for 24 h. Apoptosis was evaluated using Hoechst 33342 and propidium iodide staining. (**c**,**d**) INS-1E were transfected with siCTRL or with a siRNA targeting GPER (si*Gper1*). Cells were treated with vehicle (white bars) or G1 100 nM (blue bars) for 24 h. (**c**) Protein expression was measured by western blot. Representative images of four independent experiments are shown. (**d**) Apoptosis was evaluated using Hoechst 33342 and propidium iodide staining. (**e-h**) INS-1E (**e**,**g**) and EndoC-βH1 cells (**f**,**h**) were pre-treated with vehicle, ERα antagonist MPP 100 nM (**e**,**f**) or ERβ antagonist PHTPP 1 µM (**g**,**h**) for 3 h. Afterwards, cells were treated with vehicle (white bars) or G1 100 nM (blue bars) in the absence or presence of MPP 100 nM (**e**,**f**) or PHTPP 1 µM (**g**,**h**) for 24 h. Apoptosis was evaluated using Hoechst 33342 and propidium iodide staining. (**g**,**h**) INS-1E cells were transfected with siCTRL or siRNAs targeting ERα (si*Erα*) and ERβ (si*Erβ*) or ERα and ERβ simultaneously (si*Erα*/*Erβ*.) Cells were treated with vehicle (white bars) or G1 100 nM (blue bars) for 24 h. Apoptosis was evaluated using Hoechst 33342 and propidium iodide staining. (**e**) Dispersed islet cells from WT and BERKO mice were treated with vehicle (white bars) or G1 100 nM (blue bars) for 48 h. Apoptosis was evaluated using Hoechst 33342 and propidium iodide staining. Data are shown as means ± SEM of 4-5 independent experiments. (**a**,**b**,**e-h**) \*\**p*≤0.01 and \*\*\**p*≤0.001 vs untreated vehicle; ^###^*p*≤0.001 as indicated by bars. Two-way ANOVA. (**d**,**i**) \**p*≤0.05, \*\**p*≤0.01 and \*\*\**p*≤0.001 vs siCTRL vehicle; ^#^*p*≤0.001 as indicated by bars. Two-way ANOVA. (**j**) \*\**p*≤0.01 vs WT vehicle; ^##^*p*≤0.01 as indicated by bars. Two-way ANOVA.

### 3.4. ER heterodimerization

After ligand binding, ERα and ERβ form homodimers and heterodimers to exert their biological actions (Levin and Hammes, 2016; Razandi et al., 2004). As our data suggest that part of the BPA effect on apoptosis is directly mediated by ERα and ERβ, we sought to study if the binding of E2 or BPA would stabilize ER homo- and/or heterodimers. We performed docking and molecular dynamics simulations using human structural models of ERα and ERβ. Our models were obtained from resolved dimeric structures of the ER LBD where the protein is co-crystallized with ligands, E2 and BPA, bound to the LBD cavity that is closed by the transactivation helix H12 (Fig. 6a-c). Figure 6 depicts models of the ERα-ERα homodimer (ERαα; Fig. 6a), ERβ-ERβ homodimer (ERββ; Fig. 6b) and ERα-ERβ heterodimer (ERαβ; Fig. 6c).

**Fig. 6.**
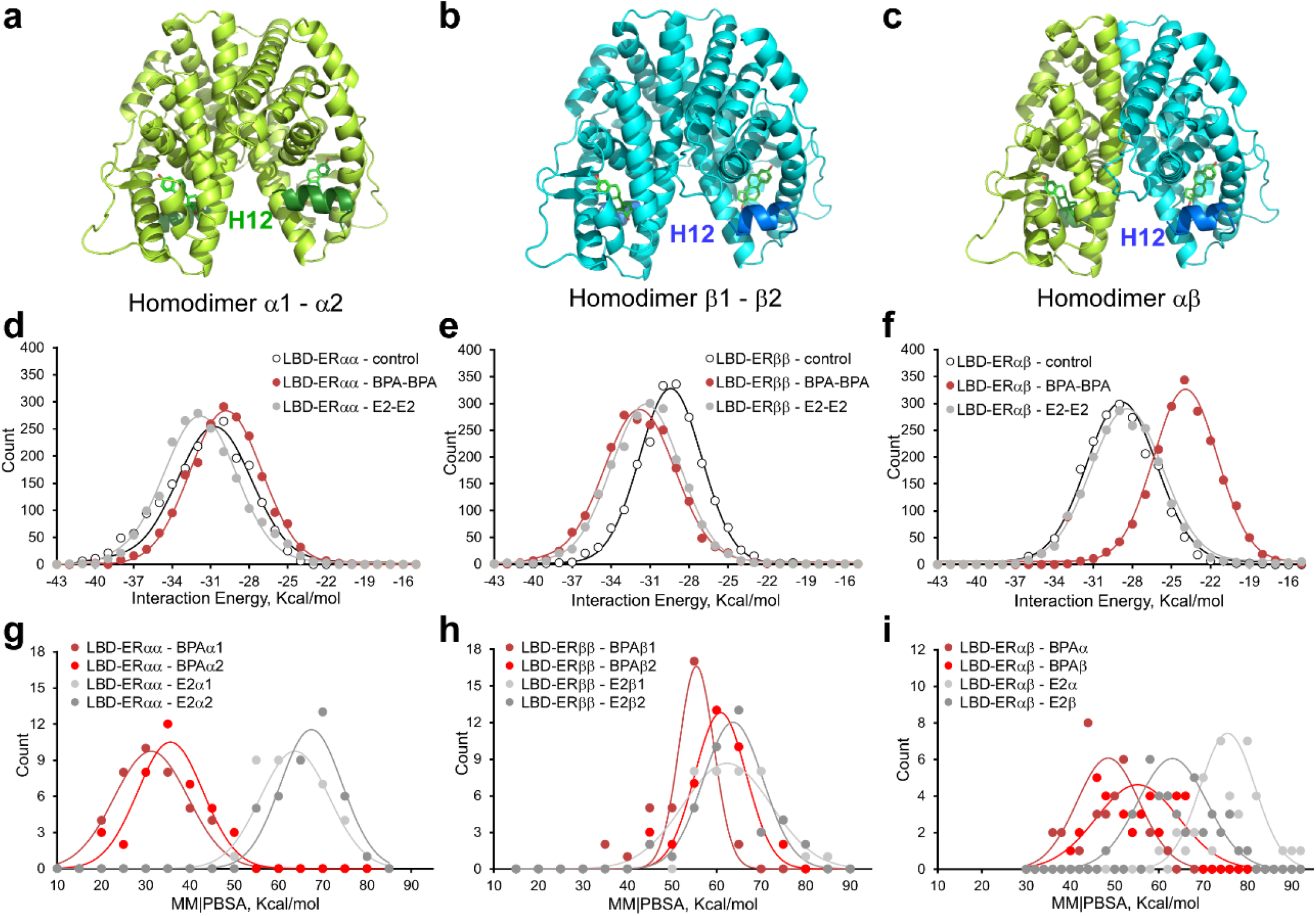
Effect of BPA and E2 on the molecular dynamics for the LBD of the homodimer-or heterodimer-ER. Secondary structure models of closed LBD-ER as (**a**) alpha homodimer, (**b**) beta homodimer and (**c**) alpha-beta heterodimer. The H12 helix has been colored dark green (alpha monomers) or dark blue (beta monomers). The structure of E2 (green sticks for carbons) is included within the LBD cavities of both subunits. (**d**) Frequency distributions of the intermolecular protein Foldx-calculated interaction energy for the subunits of the alpha homodimer, (**e**) beta homodimer and (**f**) alfa-beta heterodimer in the presence of BPA or E2 ligands in each H12 closed LBD cavity. (**g**) Frequency distributions of the molecular mechanics Poisson–Boltzmann surface area (MM/PBSA) solvation binding energy values of each ligand bound to cavity of the alpha homodimer, (**h**) beta homodimer or (**i**) alpha-beta heterodimer in the presence of BPA or E2 ligands inside of the closed LBD. YASARA software-calculated MM|PBSA values more positive energies indicate better binding of the compound bound to the protein. A Gaussian curve overlaps discrete data.

Molecular docking simulations resulted in lower Gibbs free energy changes for E2 (−11.05 and -10.86 kcal/mol for hERα-LBD and hERβ-LBD, respectively) compared with BPA (−8.15 and -8.37 kcal/mol for hERα-LBD and hERβ-LBD, respectively). These values are similar to those observed for rat ERs (Marroqui et al., 2021) and agree with the higher affinity of both receptors for E2 (compared with BPA) (Kuiper et al., 1997). We used these protein-ligand complexes as the starting point to initiate molecular dynamics simulations of the ligands within the cavity during 200 ns. Then, we analyzed the monomer-monomer interaction energy (Fig. 6d-f), the trajectory of the ligands within the LBD pocket (Supplementary Fig. 8) and the solvation binding energy of E2 and BPA bound to the LBD cavity (Fig. 6g-i).

When we examined the monomer-monomer interaction energy with E2 or BPA bound to both cavities of the LBD, we found that, for ERαα homodimers, the frequency distribution of the intermolecular interaction energy between monomers was lower for E2 than for BPA (Fig. 6d). This indicates that E2 stabilizes ERαα homodimer more than BPA. For ERββ homodimers, E2 and BPA presented a similar interaction energy, suggesting a comparable ERββ stabilization with both ligands (Fig. 6e). When the interaction energy for ERαβ heterodimers was analyzed, E2 presented a much lower interaction energy than BPA, indicating that BPA does not stabilize heterodimers (Fig. 6f). The trajectory of E2 and BPA within the cavity behaves differently; E2 showed minimal deviations during the 200 ns simulation time for both homodimers and for the heterodimer (Supplementary Fig. 8), suggesting a high stability of the hormone within the LBD pocket in the homo- and heterodimers. Conversely, BPA shows rapid structural rearrangements for both homodimers and heterodimers (Supplementary Fig. 8), a behavior previously showed for monomers (Li et al., 2018; Marroqui et al., 2021). This suggests a less stable binding of BPA compared to E2 for the three configurations of dimers.

To investigate the binding affinity of both ligands in homo- and heterodimers we analyzed the frequency distribution of the solvation binding energy values of E2 and BPA bound to each type of dimer (Wang et al., 2016) (Fig. 6g-i). Of note, positive values indicate strong ligand-protein binding (Marroqui et al., 2021). E2 binding to both homodimer cavities, LBD-ERαα (Fig. 6g) and LBD-ERββ (Fig. 6h), shows very similar values. In the heterodimer, however, the solvation energy values were higher for E2 binding to the ERα monomer forming the heterodimer (75 kcal/mol) than for E2 binding to the ERβ monomer of the heterodimer (63 kcal/mol) (Fig. 6i). For BPA, the solvation binding energy values for homodimers were 30-35 kcal/mol for LBD-ERαα (Fig. 6g) and 55-60 kcal/mol for LBD-ERββ (Fig. 6i), which indicates that BPA showed higher affinity for ERββ than for ERαα. When we analyzed the LBD-ERαβ, the frequency distribution of the solvation energy deviates from a Gaussian distribution, especially for BPA bound to the beta subunit of the heterodimer (Fig. 6i). This suggests a much lower BPA affinity for LBD-ERαβ compared with LBD-ERββ. In summary, these simulations indicate that BPA affinity is higher for ERββ than for ERαα and ERαβ. Binding of E2 should stabilize the three forms, preferentially ERαα, while BPA would mainly stabilize ERββ, to a much less extent ERαα and does not stabilize ERαβ.

To further explore how E2, BPA and G1 affect heterodimer formation we performed in situ proximity ligand assay (PLA) and co-immunoprecipitation. The PLA analysis showed a small number of heterodimers (shown as red dots) in vehicle-treated cells, which greatly increased upon E2 exposure for 24 h (Fig. 7a, Veh and E2 panels). In the presence of BPA or G1, however, very few red dots were observed, suggesting that these compounds decreased heterodimer formation (Fig. 7a, BPA and G1 panels). Co-immunoprecipitation assay indicated that ERα directly bound to ERβ (Fig. 7b-e). In agreement with the PLA data, the interaction between ERα and ERβ was augmented by E2 and significantly reduced by BPA (Fig. 7b,c) and G1 (Fig. 7d,e). Altogether, these results suggest that E2 promotes ERαβ heterodimerization while BPA and G1 seemed to disrupt ERαβ formation.

**Fig. 7.**
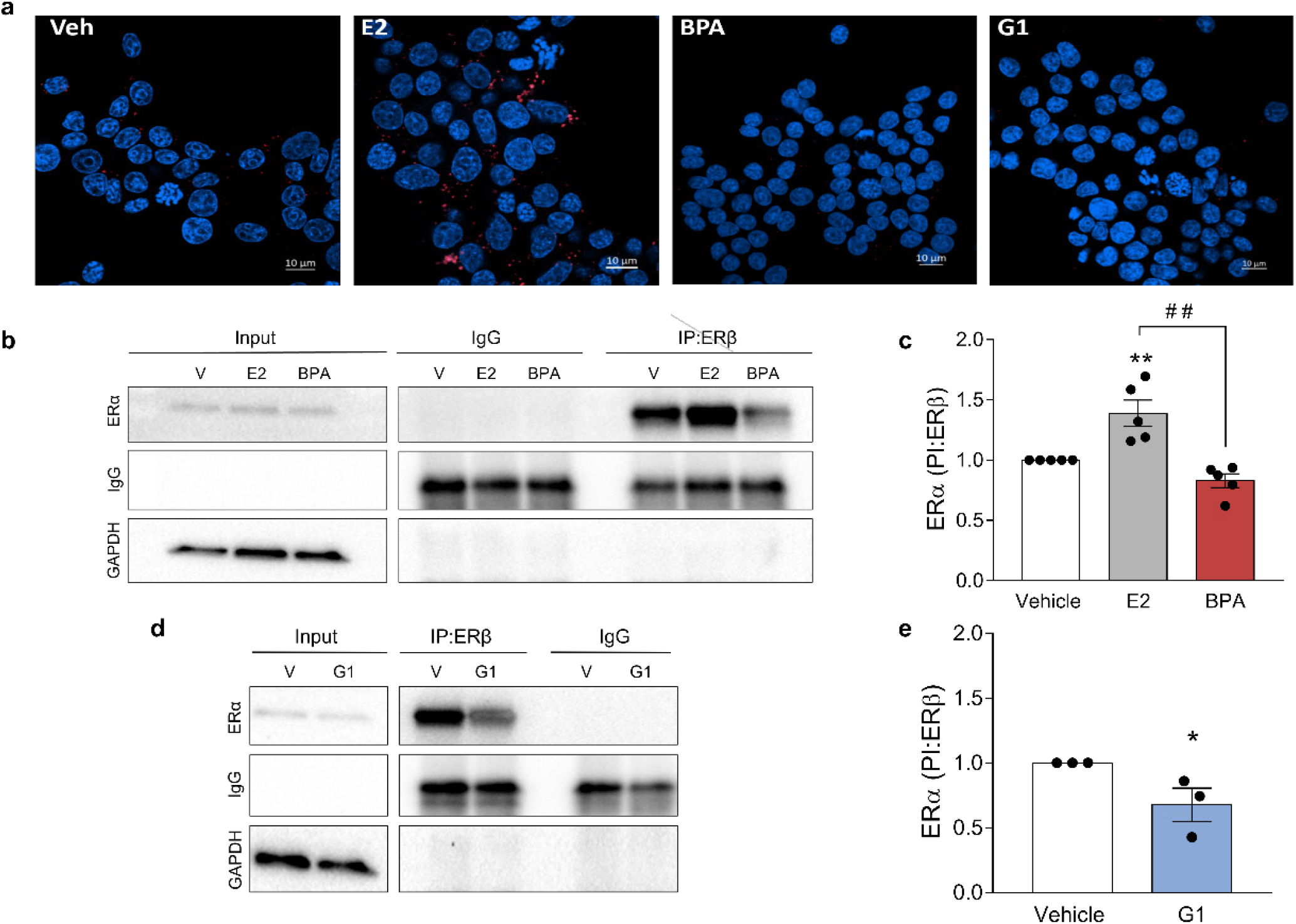
E2 and BPA have differential effects on ERα/β heterodimer formation. **(a)** INS-1E cells were treated with vehicle, E2 1 nM, BPA 1 nM or G1 100 nM for 24 h. Heterodimers were detected using in situ proximity ligand assay, where heterodimers are represented as red dots (Texas red) and nuclei are shown in blue (DAPI). (**b-e**) INS-1 E cells were treated with vehicle (white bars), E2 1 nM (grey bars), BPA 1 nM (red bars) (**b**,**c**) or G1 100 nM (blue bars) (**d**,**e**) for 24 h. Cells were lysed and proteins collected for co-immunoprecipitation with anti-ERβ antibody. Nonspecific rabbit IgG was used as a negative control. Immunoprecipitates and total protein (Input) expression were measured by western blot. Representative images of three to five independent experiments are shown (**b**,**d**) and densitometry results are presented for ERα-specific binding to ERβ (**c**,**e**). Values were normalised by GAPDH and then by the value of vehicle-treated cells of each experiment (considered as 1). Data are shown as means ± SEM of 3-5 independent experiments. (**c**) \*\**p*≤0.01 vs vehicle; ^##^*p*≤0.01 as indicated by bars. One-way ANOVA. (**e**) \**p*≤0.05 vs vehicle, by two-tailed Student’s *t* test.

## 4. Discussion

BPA is an endocrine disruptor with diabetogenic properties (Alonso-Magdalena et al., 2011; Sargis and Simmons, 2019). In adult mice, BPA increases serum insulin levels and induces insulin resistance (Alonso-Magdalena et al., 2006). When administered to pregnant mice, BPA altered insulin release, beta cell mass and the metabolome in male offspring (Alonso-Magdalena et al., 2010; Bansal et al., 2017; Cabaton et al., 2013) and alters glucose homeostasis in mothers later in life (Alonso-Magdalena et al., 2015, 2010). In humans, BPA rapidly changed plasma insulin levels (Stahlhut et al., 2018) and epidemiological studies have linked BPA exposure to type 2 diabetes development (Rancière et al., 2019; Wang et al., 2019). Here we show that environmentally relevant doses of BPA increase beta cell apoptosis in an ERα-, ERβ- and GPER-dependent manner. Moreover, molecular dynamic simulations together with PLA and co-immunoprecipitation data indicate that BPA binding to ERα and ERβ as well as GPER activation decreased ERαβ heterodimers.

It has been shown that E2 protects beta cells from several pro-apoptotic stimuli, such oxidative stress (le May et al., 2006), proinflammatory cytokines (Contreras et al., 2002) and high glucose (Kooptiwut et al., 2018). These studies suggest that ERα is crucial for pancreatic islet survival. For instance, E2 prevented streptozotocin-induced beta cell apoptosis in vivo and protected mice from insulin-deficient diabetes in an ERα-dependent manner (le May et al., 2006). Furthermore, ERα preserved mitochondrial function and attenuated endoplasmic reticulum stress in a context where ERα silencing promoted ROS production and induced beta cell apoptosis (Zhou et al., 2018). Here, ERα knockdown or treatment with the ERα antagonist MPP induced beta cell apoptosis, reinforcing an antiapoptotic role of this receptor even in the absence of added ligands. In the presence of ER ligands, we observed two different scenarios: while an ERα agonist had no effect on viability, BPA induced ROS production and beta cell apoptosis, indicating that BPA disrupts ERα antiapoptotic role. Our findings indicate that ERβ also plays a pro-survival role in beta cells, as either its silencing or antagonism with PHTPP leads to apoptosis. Interestingly, the percentage of apoptosis in ERβ-silenced cells were much higher than in ERα-deficient cells, suggesting that ERβ might be more important for beta cell survival. BPA effect on apoptosis was abrogated in ERβ-inhibited cells, indicating that, as for ERα, ERβ is also involved in BPA-induced apoptosis. The antiapoptotic protection conferred by ERs is in line with previous findings in islets from ERα and ERβ knockout mice (Liu et al., 2009).

After ligand binding, ERα and ERβ form homo- and heterodimers to perform their biological effects (Levin and Hammes, 2016). Different ER ligands stabilize ERα and ERβ homo- and heterodimers in a variety of combinations that mediate the effect of those ER ligands (Paulmurugan et al., 2011). Our bioinformatic results predicted that E2 favors the formation of ERαα, ERββ and ERαβ. BPA, however, stabilized ERββ but did not stabilize ERαβ heterodimers. Based on these results we proposed a model where, in basal conditions, ERαα, ERββ and ERαβ are antiapoptotic. Upon binding, E2 would favor ERαβ heterodimers as well as ERαα and ERββ homodimers, thus protecting beta cells from apoptosis. Conversely, ERαβ heterodimer formation does not occur in the presence of BPA; in fact, the number of ERαβ heterodimers is even decreased by BPA exposure. Therefore, as one of the three ER branches promoting survival is missing, there is an increase in apoptosis (Fig. 8).

**Fig. 8.**
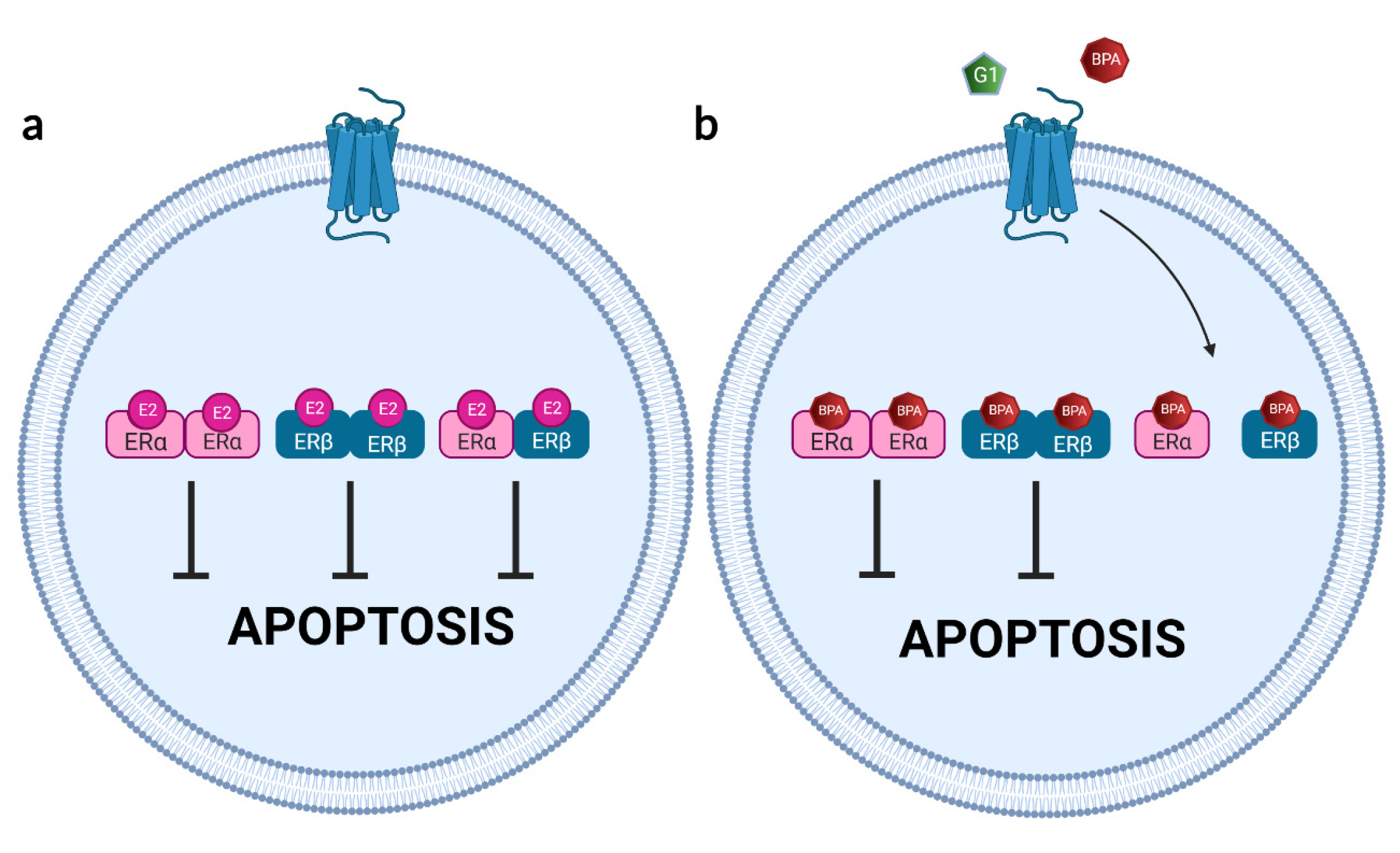
Scheme summarizing our model for E2 and BPA effects on beta cell apoptosis. (**a**) In basal conditions, ERα, ERβ homodimers and ERαβ heterodimers exist and protect beta cells from apoptosis. These dimers are also stabilized by E2. (**b**) Activation of GPER by G1 or BPA decreases ERαβ heterodimers and increases apoptosis. Binding of BPA to ERα and/or ERβ disfavors ERαβ formation. Created with BioRender.com.

A direct effect of BPA on ERα and ERβ heterodimerization might explain the part of the BPA apoptotic effect that is neither blocked by GPER antagonism nor fully achieved by GPER activation. Nonetheless, we demonstrated that GPER activation by either BPA or G1 contributes to beta cell apoptosis. As BPA binds to GPER in other cell systems (Thomas and Dong, 2006) and induces rapid nongenomic effects (Chevalier et al., 2014), it is plausible to assume that BPA acts as a GPER agonist in beta cells. As G1 and BPA effects were abolished in the presence of an ER antagonist, after ER knockdown and in islets cells from BERKO mice, we suggest that ERα and ERβ are responsible for the effect of GPER activation. A crosstalk between GPER and ERα has been described for E2 and environmental estrogens (Qie et al., 2021). Remarkably, E2, which also binds to and activates GPER (Revankar et al., 2005; Thomas et al., 2005), does not behave as G1 or BPA. Although these differences remain elusive, our data indicate that they might be related to the E2-induced increase in ERαβ heterodimer formation (Fig. 8).

Thus, along with BPA effects on insulin secretion and insulin sensitivity, BPA-induced apoptosis described by us (present data) and others (Carchia et al., 2015; Lin et al., 2013) might contribute to elucidate the diabetogenic effects of BPA described above. In the present work, BPA induces apoptosis at doses within the pico- and nanomolar range, which agrees with both *in vivo* and *in vitro* studies showing that BPA have effects at low doses (Soto et al., 2021; vom Saal and Vandenberg, 2021). Although BPA shows a low affinity for ERα and ERβ (Kuiper et al., 1998; Molina-Molina et al., 2013) and binds to GPER with an IC_50_ of 630 nM (Thomas and Dong, 2006), it is plausible to expect that an amplification of the response to BPA caused by the crosstalk among the three ERs contributes to the proapoptotic effect induced by low concentrations of BPA.

Currently, there is a lack of methods for testing EDCs that disrupt metabolism and metabolic functions (Legler et al., 2020). The MIE triggered by BPA and the subsequent increase in ROS production should help to develop novel cellular testing methods to identify other EDCs with diabetogenic properties. Moreover, our results reinforce the need to incorporate ERα and ERβ homodimer and heterodimer formation in the tests to identify EDCs, as it has been already proposed for ERα dimer formation (Kim et al., 2021). Finally, the MIE described herein reflects the complexity of the molecular mechanisms underlying EDC actions. Such complexity complicates the prediction of a given EDC effect without having exact knowledge of the MIE involved. To acquire this knowledge is a long-term goal and, therefore, a deep understanding of the MIEs should not be a strict requirement to create public health policies on EDCs (Soto and Sonnenschein, 2018).

## 5. Conclusions

The three estrogen receptors, ERα, ERβ and GPER, protect beta cells from apoptosis in basal conditions. Low doses of BPA increased beta cell apoptosis in three different cell models, namely dispersed mouse islet cells, the rat beta cell line INS-1E, and the human beta cell line EndoC-βH1. Our data indicate that EndoC-βH1 cells work as a proper human cell model for the identification of EDCs with diabetogenic activity.

The pro-apoptotic effect of BPA involves a crosstalk among the three estrogen receptors. Activation of GPER by G1 or BPA induced apoptosis via ERα and ERβ. Our results indicate that BPA directly decreases ERαβ heterodimers via ERα and ERβ or after activation of GPER. This decrease in ERαβ heterodimers disrupts the active antiapoptotic effect of ERα and ERβ in beta cells. Our results should help to develop new test methods to identify diabetogenic EDCs based on the use of human EndoC-βH1 cells, GPER activation, ROS production, and homo- and heterodimerization of ERα and ERβ.

## Abbreviations

BERKO: Estrogen Receptor β knockout
BPA: Bisphenol-A
DCF: 2’,7’-Dichlorofluorescein diacetate
E2: 17β-estradiol
ER: Estrogen receptor
ERα: Estrogen receptor α
ERβ: Estrogen receptor β
GPER: G protein-coupled estrogen receptor
LBD: Ligand binding domain
MPP: Methylpiperidinopyrazole
PHTPP: 4-(2-phenyl-5,7-bis(trifluoromethyl)pyrazolo[1,5-a]pyrimidin-3-yl)phenol
PLA: Proximity ligand assay
ROS: Reactive oxygen species
siRNA: Small interfering RNA

## Author contributions

LM and AN conceived the study. IB-C, RSDS, RMM-G, AAP-S, J-AE and LM collected and analyzed the data. IB-C, RSDS, RMM-G, AAP-S, J-AE, JM-P, LM and AN interpreted the data. RSDS, JM-P and AN drafted the manuscript. AN and LM supervised the study. RSDS, JM-P, J-AG critically revised the manuscript for important intellectual content. J-AG, J-AE, LM, JM-P and AN acquired the funding. All authors reviewed and approved the final version of the manuscript, and gave consent to publication. LM and AN are responsible for the integrity of the work as a whole.

## Acknowledgements

The authors thank Maria Luisa Navarro, Salomé Ramon, and Beatriz Bonmati Botella for their excellent technical assistance. We are grateful to the Cluster of Scientific Computing (http://ccc.umh.es/) of the Universidad Miguel Hernández de Elche for providing computing facilities.

