## Supplemental Material for "G protein-coupled estrogen receptor activation by Bisphenol-A disrupts protection from apoptosis conferred by estrogen receptors ERα and ERβ in pancreatic beta cells"

**Supplementary Table 1. List of siRNAs used in this study.**

| Code | Name | Distributor | Sequence (5'to 3') |
|---|---|---|---|
| siCTRL | Allstars Negative Control siRNA | Qiagen, Venlo, Netherlands | Sequence not provided |
| Rat si*Gper1* | Rn_Gpr30_1 FlexiTube siRNA (SI01518335) | Qiagen, Venlo, Netherlands | Sequence not provided |
| Rat si*Gper1* | Rn_Gpr30_2 FlexiTube siRNA (SI01518342) | Qiagen, Venlo, Netherlands | Sequence not provided |
| Rat si*Erα* | Esr1RSS302814 (3_RNAI) | Invitrogen, Pasley, UK | GCUACAAACCAAUGCACCAUCGAUA |
| Rat si*Erα* | Esr1RSS302815 (3_RNAI) | Invitrogen, Pasley, UK | GCUUAAUUCUGGAGUGUACACAUUU |
| Rat si*Erβ* | Esr2RSS303096 | Invitrogen, Pasley, UK | CCCAAAUGUGCUAUGGCCAACUUCU |
| Rat si*Erβ* | Esr2RSS303097 | Invitrogen, Pasley, UK | GCGUAGAAGGGAUUCUGGAAAUCUU |
| Human si*GPER1* | Hs_GPR30_1 FlexiTube siRNA (SI00430360) | Qiagen, Venlo, Netherlands | Sequence not provided |
| Human si*GPER1* | Hs_GPR30_1 FlexiTube siRNA (SI00430367) | Qiagen, Venlo, Netherlands | Sequence not provided |
| Human si*ERα* | HS ESR1 8 (SI02784101) | Qiagen, Venlo, Netherlands | Sequence not provided |
| Human si*ERβ* | HS ESR2 6 (SI03083269) | Qiagen, Venlo, Netherlands | Sequence not provided |

**Supplementary Table 2. List of primers used in this study.**

| Gene | Forward | Reverse |
|---|---|---|
| | Sequence (5'-3') | Sequence (5'-3') |
| Rat *Esr1* | CTGACAATCGACGCCAGAA | TCGTTACACACAGCACAGTAG |
| Rat *Esr2* | TGGTCATGTGAAGGATGTAAGG | TTACGCCGGTTCTTGTCTATG |
| Rat *Gper1* | TCTACACCATCTTCCTCTTCCC | ACAGGTCTGGGATAGTCATCTT |
| Rat *Gapdh* | AGTTCAACGGCACAGTCAAG | TACTCAGCACCAGCATCACC |
| Human *ESR1* | CAGATGGTCAGTGCCTTGTT | GTTGGTCAGTAAGCCCATCAT |
| Human *ESR2* | TGGGCACCTTTCTCCTTTAG | AGGTGTGTTCTAGCGATCTTG |
| Human *GPER1* | GTCTCTAAACTGCGGTCAGATG | AGCAATTCTGTGTGAGGAGTG |
| Human β-actin | CTGTACGCCAACACAGTGCT | GCTCAGGAGGAGCAATGATC |

**Supplementary Table 3. List of antibodies used in this study.**

| Target antigen | Antibody Name | Manufacturer and catalogue number (Cat no.) | Species raised in | Dilution | RRID |
|---|---|---|---|---|---|
| ERα | Estrogen Receptor alpha Monoclonal Antibody | Invitrogen; Cat no. MA5-13191 | Mouse, monoclonal | 1:1000 (WB) | AB_10986080 |
| ERα | Estrogen Receptor alpha Monoclonal Antibody (SP1) | Thermo Fisher Scientific; Cat no. MA5-14501 | Rabbit, monoclonal | 1:200 (PLA) | AB_10981779 |
| ERβ | Estrogen Receptor Beta Monoclonal Antibody | Invitrogen; Cat no. MA5-24807 | Mouse, monoclonal | 1:2000 (WB) and 2 mg/ml (IP) | AB_2717280 |
| ERβ | Estrogen Receptor beta Monoclonal Antibody (14C8) | Thermo Fisher Scientific; Cat no. MA1-23217 | Mouse, monoclonal | 1:2000 (PLA) | AB_558839 |
| GPER | Anti-G-protein coupled receptor 30 antibody | Abcam; Cat no. Ab-39742 | Rabbit, polyclonal | 1:1000 (WB) | AB_1141090 |
| α-Tubulin | Monoclonal Anti-αTubulin antibody | Sigma; Cat no. T9026 | Mouse, monoclonal | 1:5000 (WB) | AB_477593 |
| GAPDH | GAPDH (D16H11) XP Rabbit mAb antibody | Cell Signaling Technology; Cat no. 5174 | Rabbit, monoclonal | 1:1000 (WB) | AB_10622025 |
| β-Actin | β-Actin (D6A8) Rabbit mAb antibody | Cell Signaling Technology; Cat no. 8457 | Rabbit, monoclonal | 1:1000 (WB) | AB_10950489 |
| Goat anti-mouse IgG | Goat Anti-Mouse IgG (H+L) HRP Conjugate antibody | Bio-rad; Cat no. 170-6516 | Goat, Polyclonal | 1:5000 (WB) | AB_11125547 |

| | | | | | |
|---|---|---|---|---|---|
| Goat anti-rabbit IgG | Goat Anti-Rabbit IgG (H+L) HRP Conjugate antibody | Bio-rad; Cat no. 170-6515 | Goat, Polyclonal | 1:5000 (WB) | AB_11125142 |
| Goat anti-rabbit IgG | Normal Rabbit IgG antibody | Cell Signaling Technology; Cat no. 2729 | Goat, Polyclonal | Used for IP | AB_1031062 |

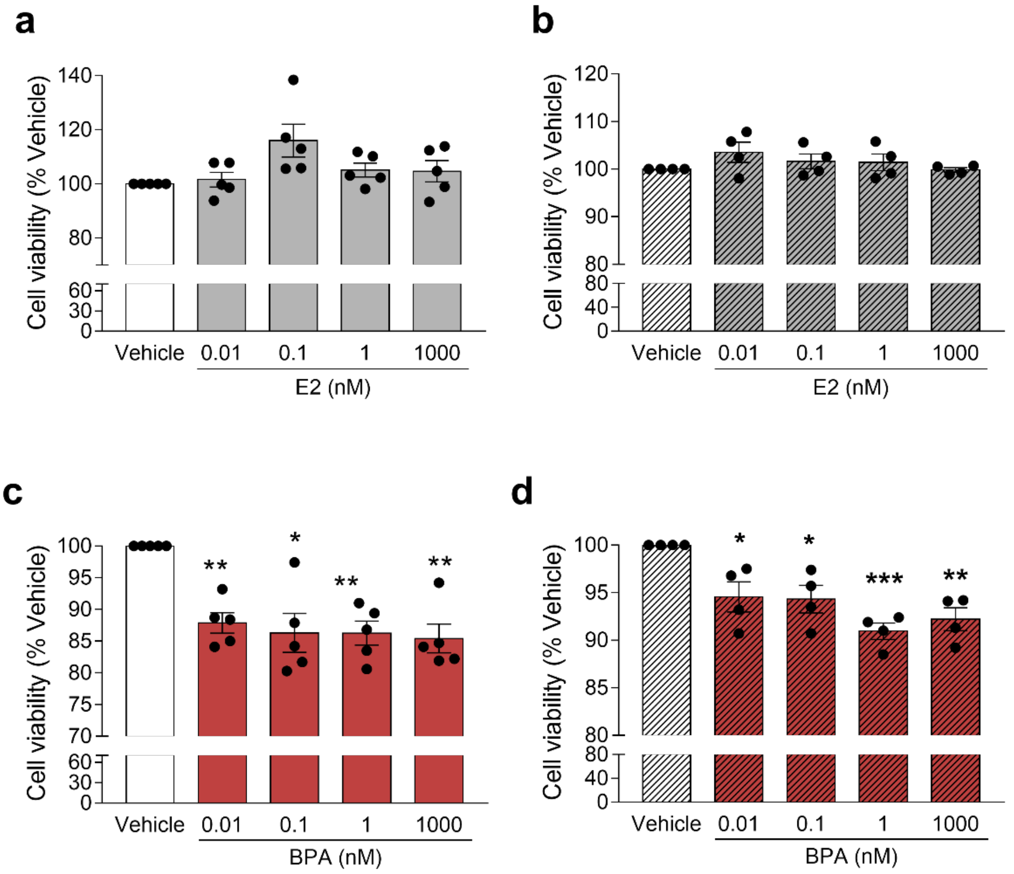


**Supplementary Fig. 1 E2 and BPA have different effects on beta cell viability.** INS-1E (**a,c**) and EndoC-βH1 cells (**b,d**) were treated with vehicle (white bars), E2 (grey bars) or BPA (red bars) for 48 h. Cell viability was evaluated by MTT assay. Data are shown as means ± SEM of 5 independent experiments. **p*≤0.05, ***p*≤0.01 and ****p*≤0.001 vs Vehicle. One-way ANOVA.

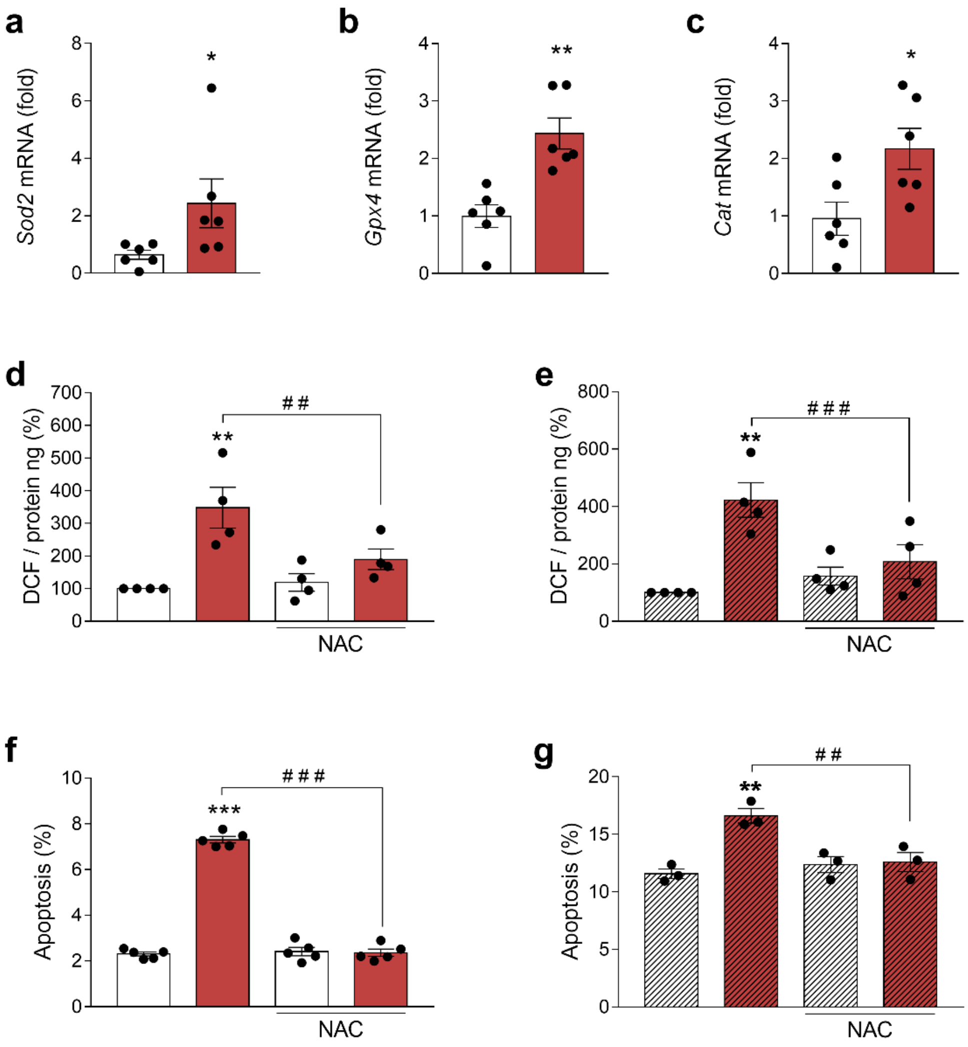


**Supplementary Fig. 2 Involvement of oxidative stress in BPA-induced apoptosis**. (**a-c**) mRNA expression of *Sod2* (**a**), *Gpx4* (**b**), and *Cat* (**c**) in INS-1E cells treated with vehicle (white bars) or BPA 1 nmol/l (red bars) for 24 h. mRNA expression was measured by qRT-PCR and normalised to the housekeeping gene *Gapdh,* and it is shown as fold vs vehicle. INS-1E (**d**) and EndoC-βH1 (**e**) cells were treated with vehicle (white bars) or BPA 1 nmol/l (red bars) in the absence or presence of N-acetylcysteine (NAC; 3 mmol/l) for 24 h. Oxidative stress was measured by oxidation of the fluorescent probe DCF and normalized by total protein. INS-1E (**f**) and EndoC-βH1 (**g**) cells were treated with vehicle (white bars) or BPA 1 nmol/l (red bars) in the absence or presence of NAC 3 mM for 24 h. Apoptosis was evaluated using Hoechst 33342 and propidium iodide staining. Data are shown as means ± SEM of 3-6 independent experiments. (**a-c**) *$p \leq 0.05$ and **$p \leq 0.01$, by two-tailed Student's *t* test. (**d-g**) **$p \leq 0.01$ and ***$p \leq 0.001$ vs the respective vehicle; $^{\#\#}p \leq 0.01$ and $^{\#\#\#}p \leq 0.001$ as indicated by bars. Two-way ANOVA.

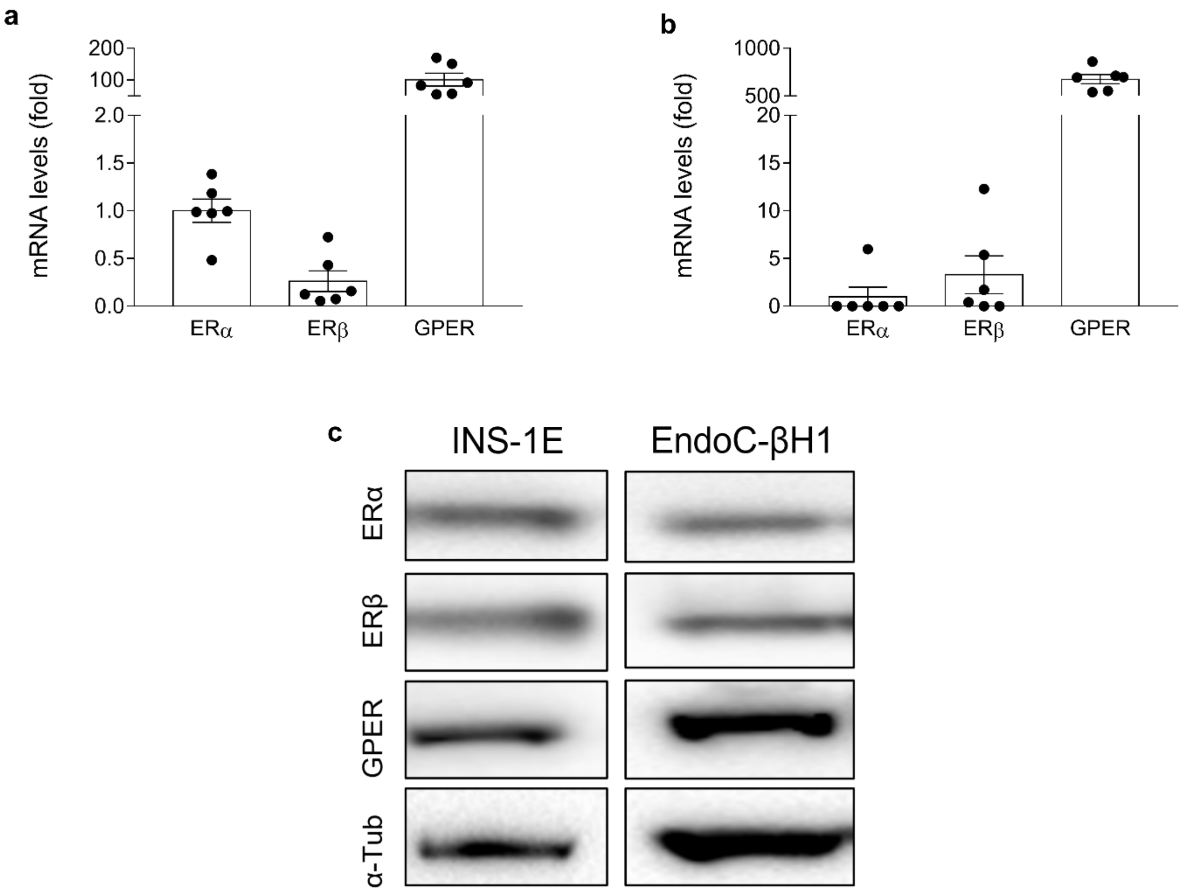


**Supplementary Fig. 3 ERα, ERβ and GPER expression in INS-1E and EndoC-βH1 cells.** mRNA expression of ERα, ERβ and GPER in INS-1E (**a**) and EndoC-βH1 cells (**b**). mRNA expression was measured by qRT-PCR and normalised to the housekeeping genes *Gapdh* (**a**) and β-actin (**b**). Data are shown as fold-change of ERα expression (considered as 1). (**c**) Protein expression of ERα, ERβ and GPER in INS-1E (left panel) and EndoC-βH1 cells (right panel) was measured by western blot.

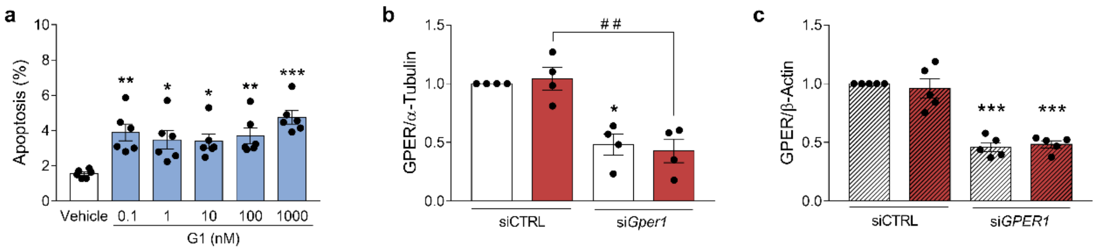


**Supplementary Fig. 4 GPER activation induces apoptosis.** (**a**) INS-1E cells were treated with vehicle (white bars) or GPER agonist G1 (blue bars) for 24 h. Apoptosis was evaluated using Hoechst 33342 and propidium iodide staining. (**b,c**) Densitometry analysis of immunoblots shown in Fig. 2e (**b**) and Fig. 2h (**c**). Values were normalised by α-tubulin (**b**) or β-actin (**c**) and then by the value of siCTRL-transfected vehicle-treated cells of each experiment (considered as 1). Data are shown as means ± SEM of 4-6 independent experiments. (**a**) *$p \leq 0.05$, **$p \leq 0.01$ and ***$p \leq 0.001$ vs Vehicle, by one-way ANOVA. (**b,c**) *$p \leq 0.05$ and ***$p \leq 0.001$ vs the respective siCTRL. $^{\#\#}p \leq 0.01$ as indicated by bars. Two-way ANOVA.

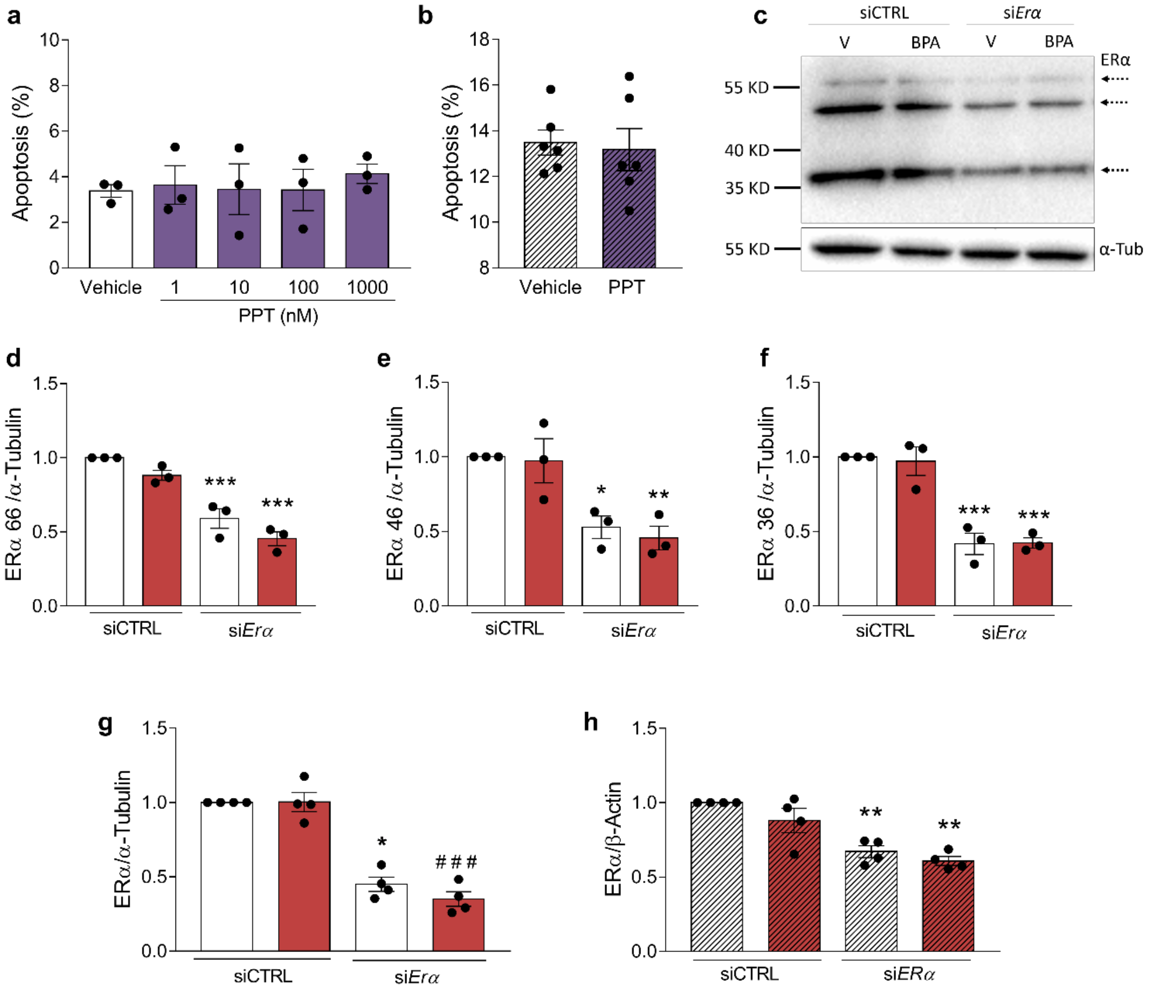


**Supplementary Fig. 5 PPT effect on viability and confirmation of ERα knockdown.** INS-1E (**a**) and EndoC-βH1 cells (**b**) were treated with vehicle or ERα agonist PPT (1 nmol/l in **b**) for 24 h. Apoptosis was evaluated using Hoechst 33342 and propidium iodide staining. (**c-f**) INS-1E cells were transfected with siCTRL or with a siRNA targeting ERα (si*Erα*). Cells were treated with vehicle (white bars) or BPA 1 nmol/l (red bars) for 24 h. Protein expression was measured by western blot. Representative images of three independent experiments are shown (**c**) and densitometry results are presented for different ERα variants, namely ERα 66 (**d**), ERα 46 (**e**) and ERα 36 (**f**). Values were normalised by α-tubulin (α-tub) and then by the value of siCTRL-transfected vehicle-treated cells of each experiment (considered as 1). (**g,h**) Densitometry analysis of immunoblots shown in Fig. 3d (**g**) and Fig. 3g (**h**). Values were normalised by α-tubulin or β-actin and then by the value of siCTRL-transfected vehicle-treated cells of each experiment (considered as 1). Data are shown as means ± SEM of 3-4 independent experiments. (**d-h**) *$p \le 0.05$, **$p \le 0.01$, ***$p \le 0.001$ vs the respective siCTRL, by two-way ANOVA. PPT, propylpyrazoletriol.

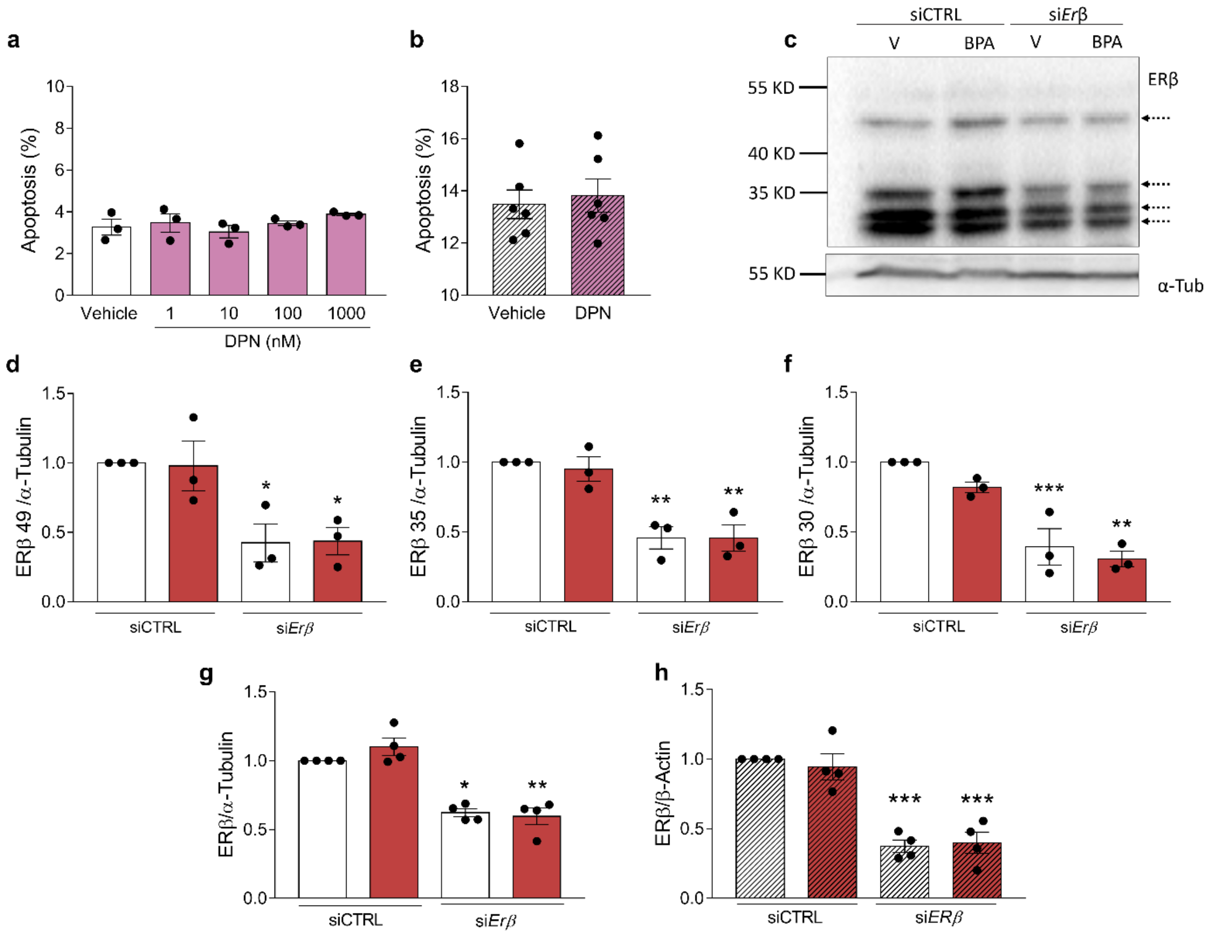


**Supplementary Fig. 6 DPN effect on viability and confirmation of ERβ knockdown.** INS-1E (**a**) and EndoC-βH1 cells (**b**) were treated with vehicle or ERβ agonist DPN (1 nmol/l in **b**) for 24 h. Apoptosis was evaluated using Hoechst 33342 and propidium iodide staining. (**c-f**) INS-1E cells were transfected with siCTRL or with a siRNA targeting ERα (si*Erβ*). Cells were treated with vehicle (white bars) or BPA 1 nmol/l (red bars) for 24 h. Protein expression was measured by western blot. Representative images of three independent experiments are shown (**c**) and densitometry results are presented for different ERβ variants, namely ERβ 49 (**d**), ERβ 35 (**e**) and ERβ 30 (**f**). Values were normalised by α-tubulin (α-tub) and then by the value of siCTRL-transfected vehicle-treated cells of each experiment (considered as 1). (**g,h**) Densitometry analysis of immunoblots shown in Fig. 4c (**g**) and Fig. 4f (**h**). Values were normalised by α-tubulin or β-actin and then by the value of siCTRL-transfected vehicle-treated cells of each experiment (considered as 1). Data are shown as means ± SEM of 3-6 independent experiments: (d-h) *$p \leq 0.05$, **$p \leq 0.01$, ***$p \leq 0.001$ vs the respective siCTRL, by two-way ANOVA. DPN, diarylpropionitrile.

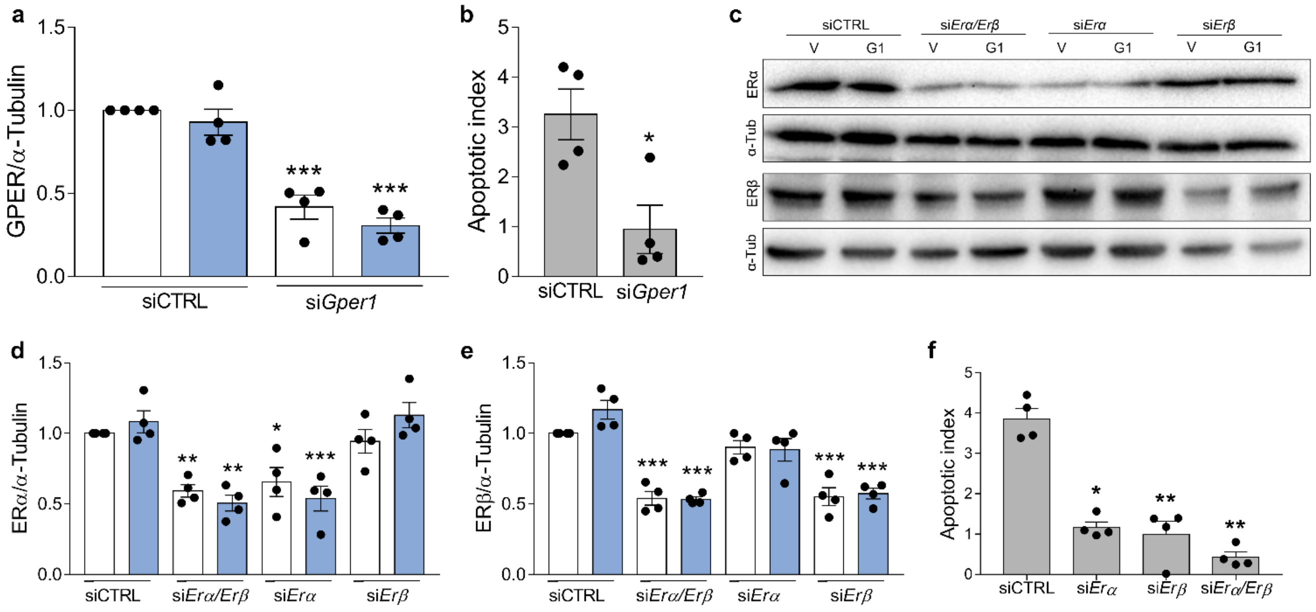


**Supplementary Fig. 7 GPER requires ERα and ERβ to induce apoptosis.** (**a**) Densitometry analysis of immunoblots shown in Fig. 5c. Values were normalised by α-tubulin and then by the value of siCTRL-transfected vehicle-treated cells of each experiment (considered as 1). (**b**) G1-induced apoptosis data from Fig. 5d are presented as apoptotic index. (**c-e**) Related to Fig. 5i. Protein expression was measured by western blot. Representative images of four independent experiments are shown (**c**) and densitometry results are presented for ERα (**d**) or ERβ (**e**). (**f**) G1-induced apoptosis data from Fig. 5i are presented as apoptotic index. Data are shown as means ± SEM of 4 independent experiments. (**a,d,e**) *$p \leq 0.05$, **$p \leq 0.01$, ***$p \leq 0.001$ vs the respective siCTRL, by two-way ANOVA. (**b**) *$p \leq 0.05$, by two-tailed Student's *t* test. (**f**) *$p \leq 0.05$ and **$p \leq 0.01$ vs siCTRL, by one-way ANOVA.

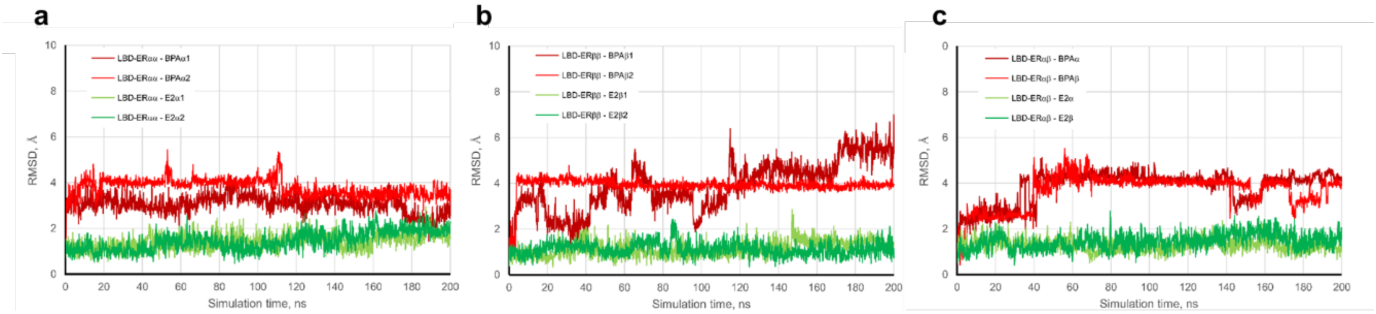


**Supplementary Fig. 8 Molecular dynamics simulation of homo- and heterodimers of hLBD-ER**. Analysis of the trajectory of the ligands bound to the closed cavity of the LBD-ER for homodimers α/α (**a**), homodimers β/β (**b**) and heterodimers α/β (**c**).
